# Somatic epimutations enable single-cell lineage tracing in native hematopoiesis across the murine and human lifespan

**DOI:** 10.1101/2024.04.01.587514

**Authors:** Michael Scherer, Indranil Singh, Martina Braun, Chelsea Szu-Tu, Michael Kardorff, Julia Rühle, Robert Frömel, Sergi Beneyto-Calabuig, Simon Raffel, Alejo Rodriguez-Fraticelli, Lars Velten

## Abstract

Current approaches to lineage tracing of stem cell clones require genetic engineering or rely on sparse somatic DNA variants, which are difficult to capture at single-cell resolution. Here, we show that targeted single-cell measurements of DNA methylation at single-CpG resolution deliver joint information about cellular differentiation state and clonal identities. We develop EPI-clone, a droplet-based method for transgene-free lineage tracing, and apply it to study hematopoiesis, capturing hundreds of clonal trajectories across almost 100,000 single-cells. Using ground-truth genetic barcodes, we demonstrate that EPI-clone accurately identifies clonal lineages throughout hematopoietic differentiation. Applied to unperturbed hematopoiesis, we describe an overall decline of clonal complexity during murine ageing and the expansion of rare low-output stem cell clones. In aged human donors, we identified expanded hematopoietic clones with and without genetic lesions, and various degrees of clonal complexity. Taken together, EPI-clone enables accurate and transgene-free single-cell lineage tracing at scale.

## Introduction

Lineage tracing by genetic or physical labels has been an important tool in developmental and stem cell biology for more than a century^1^. Genetic barcoding compatible with single-cell readouts, in particular single-cell RNA-seq, represents the most advanced lineage tracing tool available, as it provides information on the cellular output of hundreds or thousands of stem cell clones together with cell state information on the stem cell itself^2–9^. These technologies have recently generated new insights into clonal function in healthy, pre-cancerous and cancerous tissues^10–12^. However, such methods require complex genetic engineering and are therefore restricted in applications, including profiling stem cell fates in native development and in human tissues.

Recent efforts have focused on developing methods for lineage tracing that rely on endogenous clonal markers (e.g., mutations), do not require genetic engineering, and still allow tracing of many stem cell clones in parallel. Whole genome sequencing provides such a dense lineage picture thanks to the mapping of somatic DNA mutations^13,14^, but it suffers from very limited throughput and lacks information about the cell state. On the other hand, spontaneous mitochondrial DNA (mtDNA) mutations can be captured by single-cell RNA-seq or ATAC-seq^15–18^. However, a recent preprint analyzed both mtDNA and nuclear somatic mutations from clonally expanded hematopoietic stem cells, and found that most apparent mitochondrial genetic variation does not reflect the phylogenetic signal obtained from whole-genome sequencing of the nuclear genome^19^, raising doubts whether cellular phylogenies can be faithfully reconstructed with that approach.

Epimutations, defined as the spontaneous loss/gain of DNA methylation at individual CpG dinucleotides, have been explored as potential clonal labels in cancer^20–22^. The clonal signal present in DNA methylation data is stronger than in gene expression or chromatin accessibility data^23^ and clonal signals have been identified in bulk DNA methylation data^24,25^. On the other hand, DNA methylation is a highly cell-type-specific epigenetic mark^26–29^, and there is no prior evidence that it can be exploited to trace lineages of single cells in healthy tissues. Moreover, single-cell DNA methylation (scDNAm) datasets are typically generated by bisulfite sequencing and related methods, which is expensive, low-throughput, and possibly too noisy to capture epimutations at individual CpGs.

Here, we demonstrate that a targeted, high-confidence, single-cell readout of DNA methylation at single CpG resolution is sufficient to track clones in adult blood formation (hematopoiesis) and further provides detailed information on cell state. We present a novel method, EPI-Clone, that extracts both clonal and cell state information from targeted single-cell DNA methylation data. EPI-Clone builds upon scTAM-seq, which uses the commercially available Mission Bio Tapestri platform, to read out methylation of several hundred CpGs in up to tens of thousands of single cells at a time, with a dropout rate of ∼7%^30^. We applied EPI-clone to barcoded hematopoietic progenitor and mature myeloid cells, as well as in native human and mouse hematopoiesis and were able to accurately identify clone of origin for most cells. EPI-clone achieved high overlap with ground-truth genetic barcodes and uncovered alterations to the clonal composition and clonal function of hematopoietic stem cells in unperturbed ageing.

## Results

### DNA methylation states are influenced both by clonal identity and by differentiation state

To create a ground-truth dataset of clonal identity and DNA methylation, we labelled murine hematopoietic stem cells (HSCs) with lentiviral barcodes using the LARRY system^2^. Labeled HSCs were transplanted into lethally irradiated mice, allowed to reconstitute all blood populations, and profiled five months later. Sorted stem and progenitor cells from bone marrow (HSPCs, sorted as Lin-c-Kit+ [LKs] with additional enrichment of Lin-Sca-1+c-Kit+ [LSKs]) as well as mature myeloid cells (sorted by the expression of CD11b) were profiled by scTAM-seq (Figure 1a, Figure S1a, b) in five experimental batches (i) Main LARRY experiment with four mice (13,885 cells) (ii) replicate LARRY experiment with two additional mice (7,896 cells), (iii) wildtype mouse without LARRY barcodes (7,001 cells), (iv) mature myeloid cells from lung, peripheral blood (PB) and bone marrow (BM) (9,204 cells), and (v) comparison between young and aged wildtype mice (15,907 cells). We designed a panel of 453 CpGs based on bulk HSPC methylome data^28^ using differential methylation between different stem- and progenitor cell subsets^31^ (Figure 1b). Additionally, we included CpGs in our panel that are variably methylated within HSCs by leveraging read heterogeneity in bulk data^32^ (Figure S1c, d and Methods). The expression of 20 surface proteins (Table S1) was simultaneously profiled using oligo-tagged antibodies to obtain independent information on cellular differentiation.

**Figure 1:**
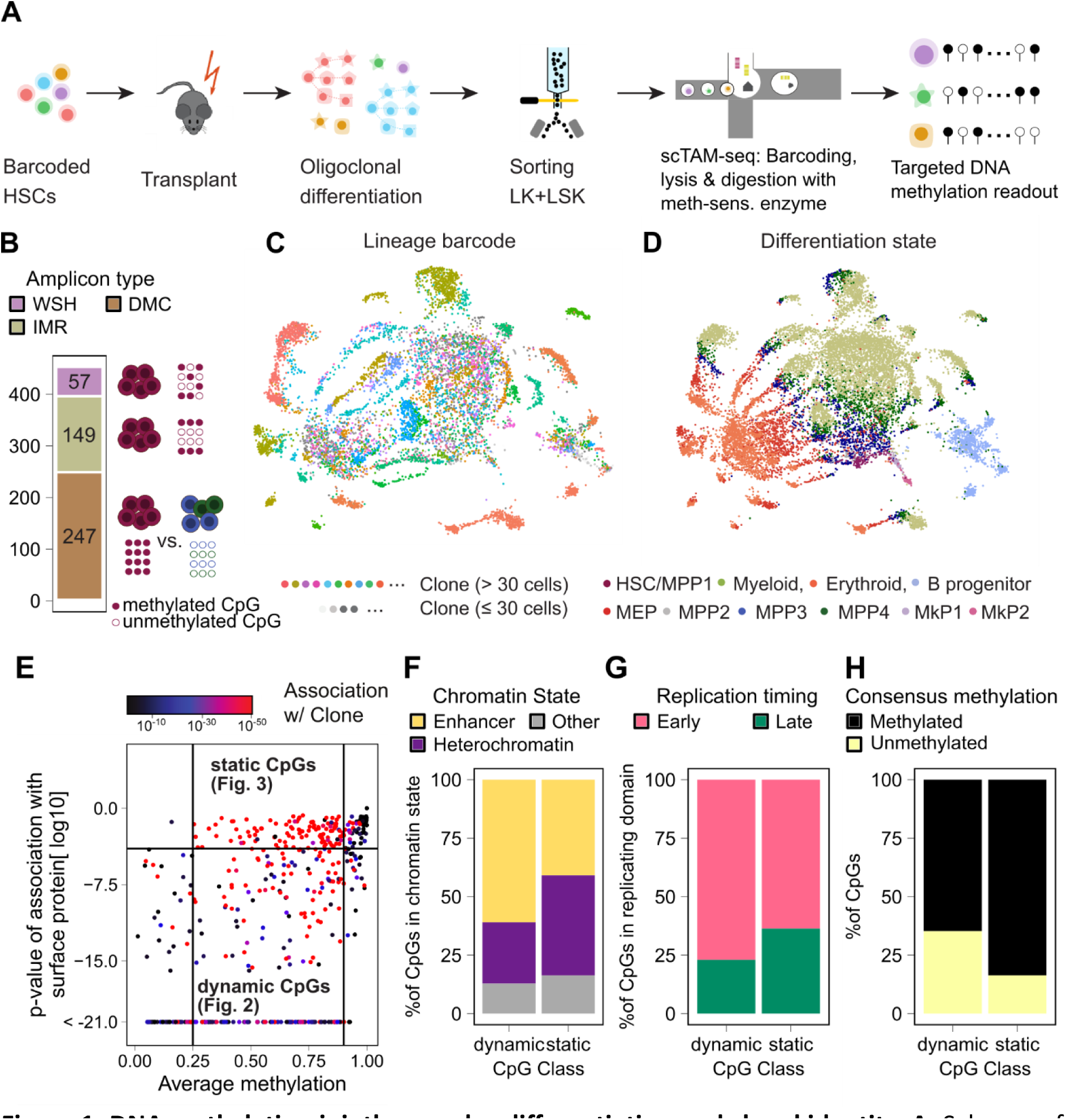
DNA methylation jointly encodes differentiation and clonal identity. **A.** Scheme of the experimental design. **B.** Overview of the 453 CpGs covered by our based on differential methylation in bulk data and performance in an undigested control experiment. WSH=within-sample heterogeneity, DMC=differentially methylated cytosine, IMR=intermediately methylated region **C.** uMAP of DNA methylation data from HSPCs from the LARRY main experiment (four mice). The color highlights clone, as defined from the LARRY barcode. **D.** Same uMAP as in C indicating cell states, see Figure 2 for how cell state was defined. **E.** Scatter plot depicting, for n=453 CpGs, the average methylation rate (x axis), the statistical association with surface protein expression (y axis, see Methods) and the statistical association with LARRY clonal labels (color coded, p value from a chi-square test). The CpGs in the upper/lower central rectangle were defined as static/dynamic CpGs, respectively. **F, G.** Bar chart depicting the fraction of static and dynamic CpGs within annotated enhancer/heterochromatic regions and within early/late replicating domains, respectively. **H.** Average methylation state across all profiled cells for the dynamic/static CpGs. CpGs with more than 50% methylated cells were termed methylated, the remaining ones unmethylated.

A uMAP display of the data from the main experiment revealed that DNA methylation jointly captures two layers of information: While cells clustered by their clonal identity defined through LARRY barcodes (Figure 1c) they also clustered by differentiation state (Figure 1d, Figure S2a, and see below for how differentiation states were annotated). We hypothesized that different subsets of CpGs might be impacted by clonal identity and differentiation, respectively. We identified differentiation associated, *dynamic* CpGs by testing for the association with the expression of any surface protein, and found that the remaining, *static* CpGs were frequently associated with clonal identity, as defined through LARRY (Figure 1e). *Dynamic* CpGs were enriched in enhancer elements, whereas the *static* CpGs were preferentially located in heterochromatic regions^33^ (Figure 1f). Additionally, static CpGs were more often located in late replicating domains^34^ as compared to dynamic CpGs (Figure 1g). Static CpGs stochastically gain/lose (preferentially lose, Figure 1h) methylation. We thus hypothesize that DNA methylation changes in static CpGs correspond to stochastic, but clonally heritable *epimutations,* which are present at the moment of LARRY barcoding, and remain largely stable through time and differentiation.

### DNA methylation delivers a high-resolution map of murine hematopoiesis

We next sought to further dissect the two layers of information in our DNA methylation landscape: differentiation state (Figure 2) and clonal identity (Figure 3). We expect epimutations to be clone- and thus batch-specific, since they occur stochastically. Indeed, we found that by integrating methylation data from 28,782 cells across three experiments profiling HSPCs (main and replicate LARRY experiment, native hematopoiesis), clonal information was effectively removed. In the resulting low dimensional embedding, we found that most variation was driven by differentiation along four trajectories (myeloid, erythroid, lymphoid and megakaryocytic differentiation, Figure 2a). A similar landscape was obtained by performing dimensionality reduction using only the *dynamic* CpGs in the main LARRY experiment (Figure S2b). Since we also performed scRNA-seq on the same samples, we could compare the DNAm uMAP with a transcriptomic uMAP (Figure 2b). We observed an overall similar topology with the four main differentiation trajectories. scRNA-seq achieved a higher resolution in the late myeloid progenitors, compared to the stem- and progenitor-cell-focused methylation panel (Figure 1b).

**Figure 2:**
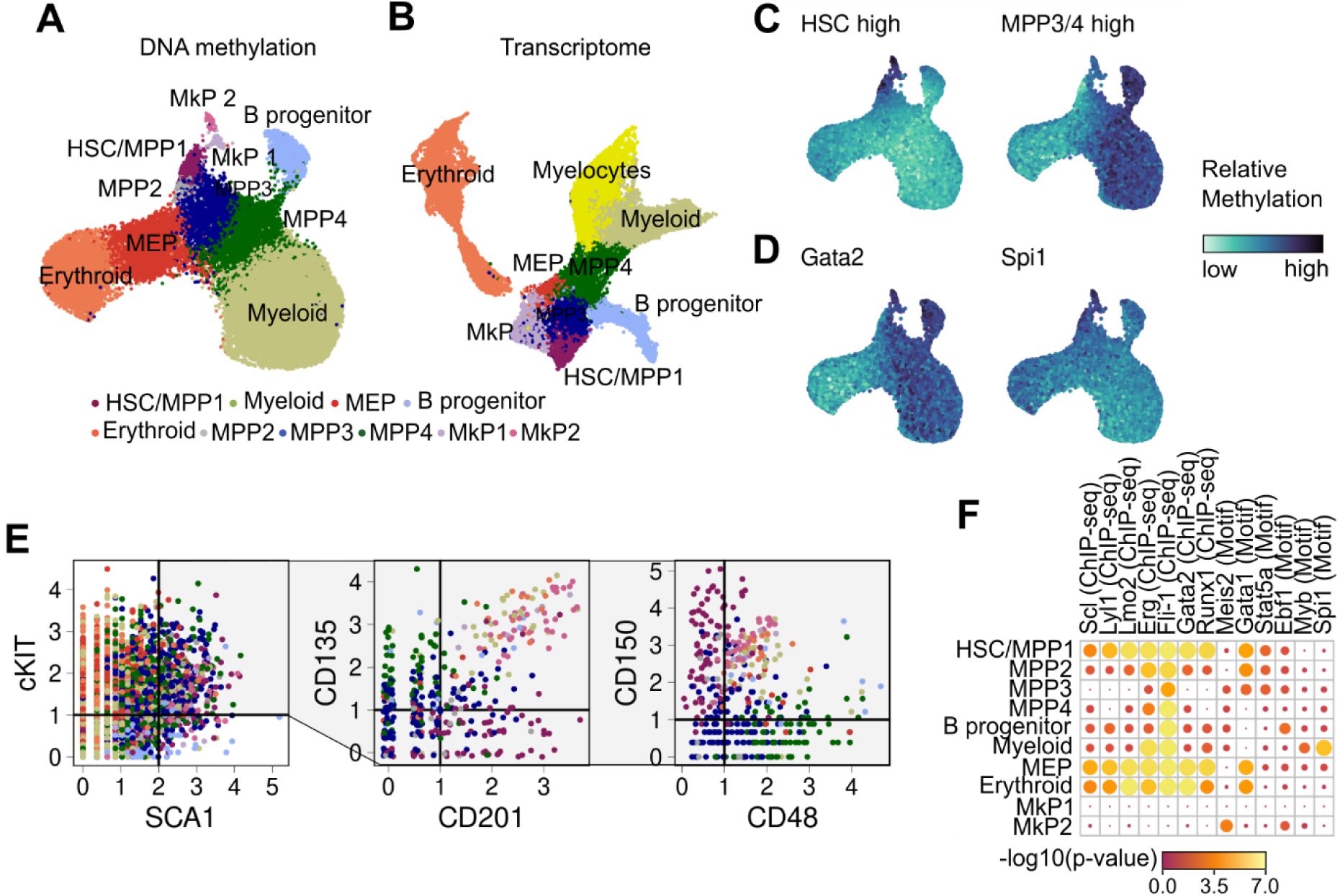
Single-cell DNA methylation profiles deliver a high-resolution map of mouse hematopoiesis. **A.** uMAP of DNA methylation data for HSPCs from the LARRY main experiment, the LARRY replicate experiment and from native hematopoiesis. Batch correction was applied prior to uMAP. Colors highlight groups identified from unsupervised clustering. **B.** uMAP of transcriptomic data from the same experiments. **C.** uMAP as in A, highlighting relative methylation state of cells across all CpGs that are methylated in HSCs or MPP3/4 in bulk data. **D.** uMAP as in A, highlighting relative methylation states of CpGs located in vicinity of TFBS. **E.** Surface protein expression of Sca1, c-Kit, CD135, CD201, CD48, and CD150. The CD135/CD201 and CD48/CD150 plots only show Lin^-^Sca-1^+^c-KIT^+^ cells and the color codes corresponds to the cell type clusters defined in the DNAm uMAP **F.** Enrichment of CpGs specifically unmethylated in a cell-type cluster according to the vicinity to the annotated TFBS.

**Figure 3:**
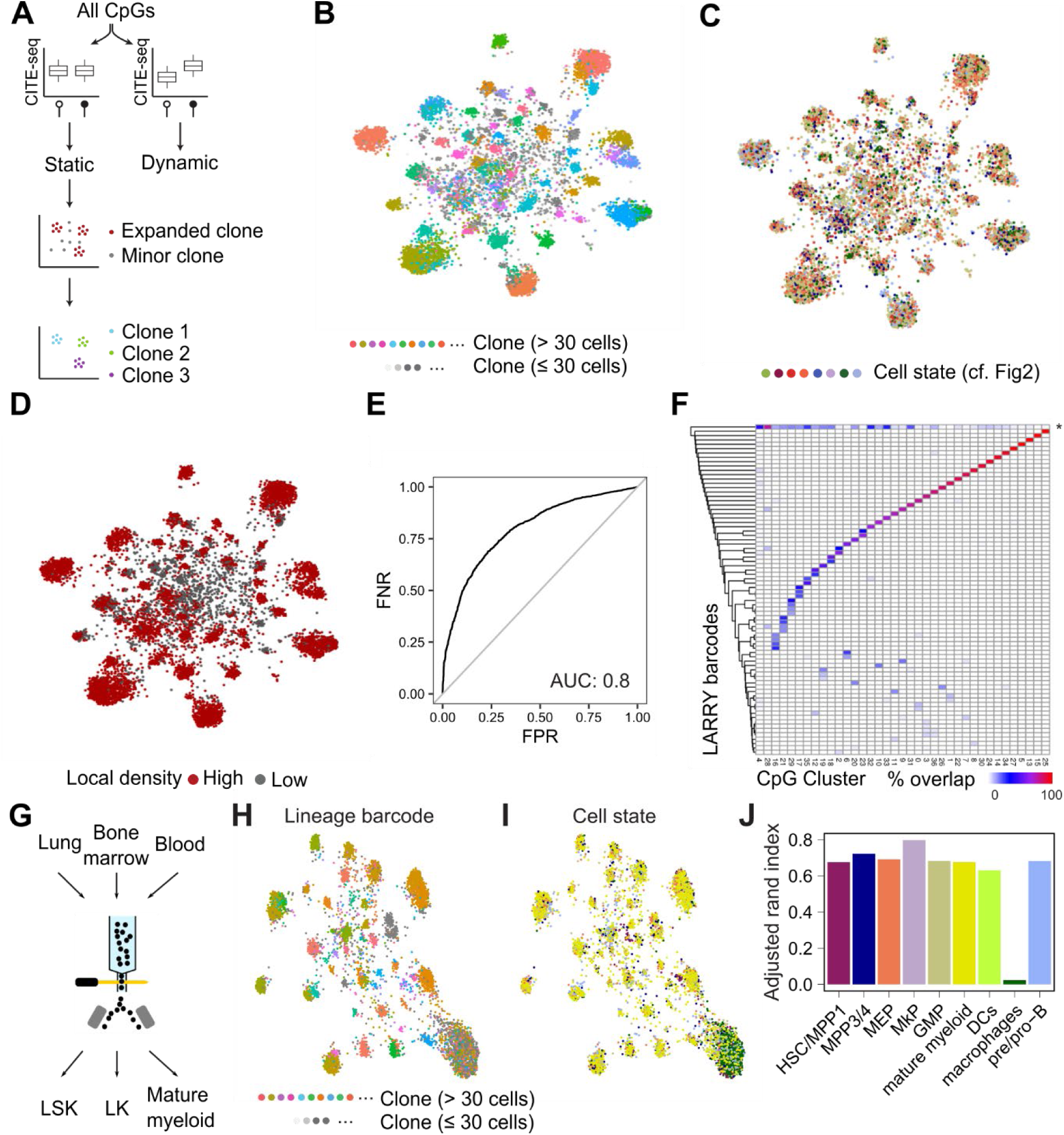
EPI-clone reliably identifies clones only from DNA methylation data. **A.** Schematic overview of EPI-clone. **B.** uMAP of DNA methylation computed on static CpGs only for the LARRY main experiment, highlighting clonal identity as defined by LARRY barcodes. Only cells carrying a LARRY barcode are shown. **C, D.** Same uMAP as in B highlighting the cells by cell state as defined in Figure 2 (C) and those that were selected as part of expanded clones based on local density in PCA space (D). **E.** Receiver-Operating Characteristics Curve visualizing the performance of classifying cells into expanded and non-expanded clones based on local density in PCA space. **F.** Heatmap depicting the association between LARRY barcode and methylation-based clonal cluster identified by EPI-Clone. The row labeled with an asterisk contains all LARRY clones smaller than 30 cells. **G**. Sorting scheme for the mature myeloid experiment. **H, I.** UMAP representation of the mature myeloid experiment. Cells cluster both by clonal identity (LARRY barcode, H) and cell state (I). **J.** Adjusted rand indices between the ground truth clonal label (LARRY) and the clones identified by EPI-Clone stratified by cell type

To annotate cell states from scDNAm data, we used three layers of information: i) bulk methylation profiles (Figure 2c, Figure S2c), ii) the relative methylation states of important lineage-specific transcription factor binding sites (TFBS, Figure 2d) and iii) the expression of surface proteins (Figure 2e, Figure S2d). We identified cell-state-specific demethylation of CpGs neighboring crucial TFBS including Gata2 (an erythroid factor), Ebf1 (lymphoid) and Spi1 (myeloid) (Figure 2d, f). scTAM-seq data revealed a cluster of HSCs and early multipotent progenitors (MPP1, also called short-term or active HSCs), several further MPP subsets (MPP2, MPP3, MPP4), myeloid, erythroid, and B cell progenitors, as well as two subsets of megakaryocyte progenitors, one of which was masked in scRNA-seq data (Supplementary Note, Figure S2e, f, g). Taken together, an analysis of *dynamic* CpGs provides a first DNA methylation-based map of murine HSC differentiation at single-CpG resolution and contains two orders of magnitude more cells than two previous, single-cell bisulfite sequencing datasets^35,36^.

### EPI-Clone detects clonal identities from DNA methylation data only

We then focused on exploiting the static CpGs to dissect clonal identity. To this end, we developed the EPI-Clone algorithm (Figure 3a). After selecting *static* CpGs based on absence of correlation with surface antigen expression, EPI-Clone performs dimensionality reduction on these CpGs only. We thereby found that expanded clones (>30 cells) marked by single LARRY barcodes clustered separately, with no influence of cell state (Figure 3b, c). By contrast, cells from small clones profiled with less than 30 cells (corresponding to a relative size of 0.2%) were interspersed between clusters (Figure 3d). EPI-Clone identifies cells that belong to expanded clones based on a higher local density in PCA space spanned by the *static* CpGs (Figure 3d). Thereby, EPI-Clone correctly identifies cells from expanded clones with an AUC of the ROC 0.8, using the LARRY clone sizes as ground truth (Figure 3e). Subsequently, EPI-Clone clusters cells from expanded clones by clonal identity, achieving an adjusted Rand Index of 0.88 relative to ground-truth clonal barcodes (Figure 3f). Quantitatively and qualitatively similar results were obtained on a biological replicate (Figure S3; AUC: 0.71, adjusted Rand Index: 0.84). Importantly, static CpG methylation remained associated with ground-truth genetic barcodes several months after the lentiviral labelling event, indicating that these epimutations are highly stable over long periods of time.

While epimutational clonal signals are stably maintained in blood stem and progenitors cells, we wondered whether EPI-Clone is also applicable to mature immune cells. To that end, we collected mature myeloid cells from lung, bone marrow and peripheral blood, as well as stem cells (LSKs) from the same animals and profiled them with scTAM-seq (Figure 3g). The mature myeloid cells clustered downstream of their progenitor cells in a low-dimensional representation spanned by the dynamic CpGs (Figure S4a-d). Using the static CpGs, EPI-Clone again yielded a clonal clustering that recapitulated ground truth clonal labels (Figure 3h, i; Figure S4e-h). Notably, we found no clone-associated clusters in tissue-resident macrophages (Figure 3h, Figure S4h). However, unlike other mature myeloid cells, the tissue-resident macrophages also carried multiple LARRY barcodes (Figure S4i), suggesting that macrophages phagocytose genetic material from other cells, thereby obscuring DNA-based lineage tracing. When removing the macrophages from the dataset, we found a high overlap between EPI-Clone clusters and LARRY barcodes with an Adjusted Rand Index of 0.75 (Figure 3i). The result on the mature myeloid cells shows that clonal information encoded in the DNA methylation state is maintained until differentiated myeloid cells, and that EPI-Clone can associate stem cells to their mature clonal progeny in tissues. Furthermore, it demonstrates that the target panel designed for HSPCs also performs well for identifying mature myeloid cell states in tissues.

### The number of blood-producing clones decreases with organism aging

Since EPI-clone can provide joint information on cell state of progenitors, clonal identity, and clonally derived progeny, it might be an ideal method to characterize the clonal dynamics of fully unperturbed hematopoiesis. In contrast to the transplantation setting, native hematopoiesis has been described as polyclonal^8,10^, where several thousand clones contribute at variable rates to blood formation. To investigate whether EPI-clone also identifies clones in native hematopoiesis, we investigated a bone marrow sample from an untreated, young mouse (12 weeks old). We found that 41.6% of cells were part of expanded clones that individually made up 1-4% of total HSPCs (Figure 4a). These clone sizes are in line with a study that genetically barcoded adult hematopoietic clones *in situ*^10^ (Figure 4b). The remaining 58.4% of cells were classified as belonging to small, non-expanded clones. In conclusion, EPI-clone proves to be an effective transgene-free method to identify hematopoietic clones *in situ*.

**Figure 4:**
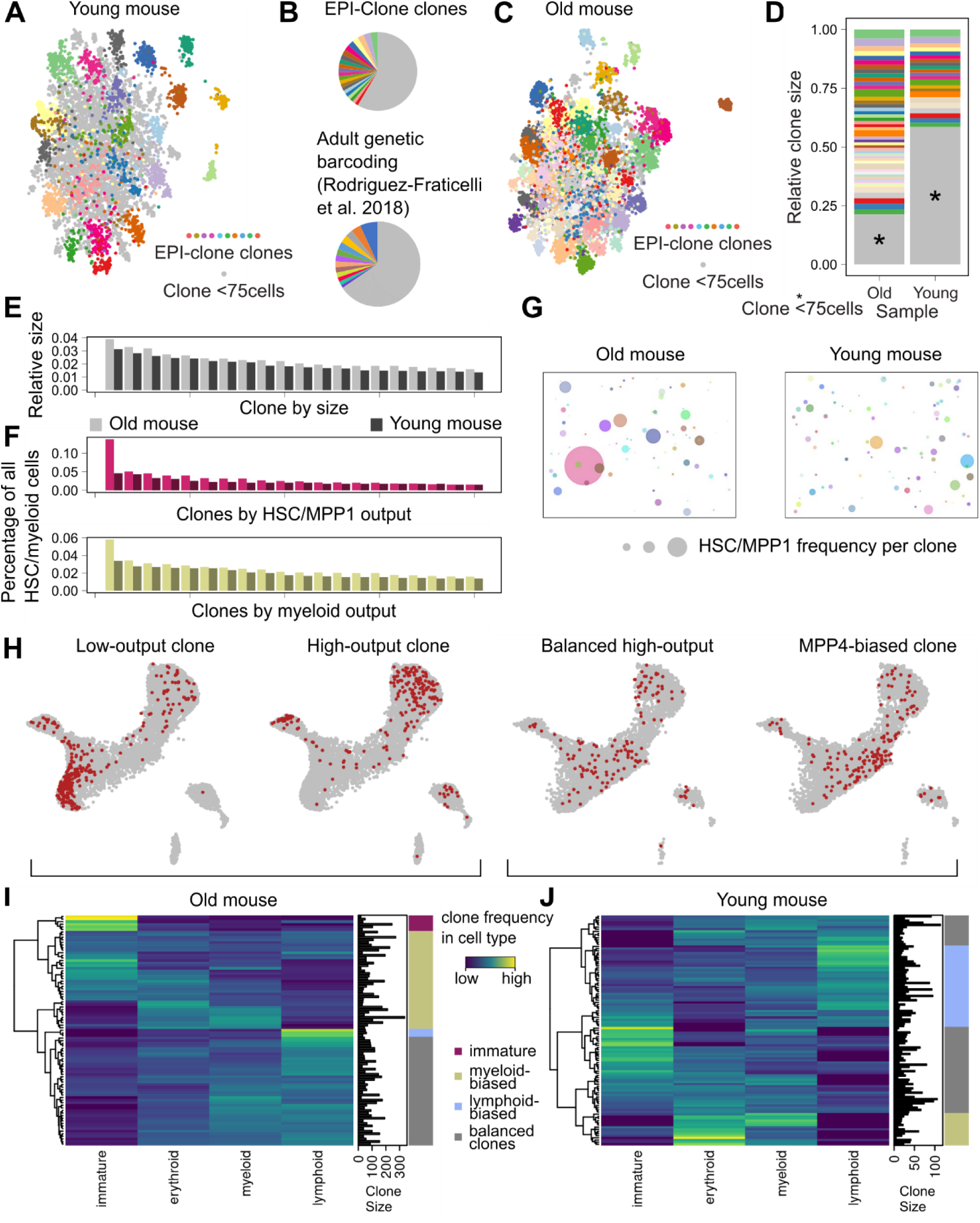
Blood production in age is characterized by a decrease in clonal complexity. **A.** UMAP based on the static CpGs for a native, young mouse. **B.** Pie charts depicting size of expanded clones identified in this study and in a study where cells were labeled using a DNA barcode in the adult^10^. In the EPI-clone pie chart, grey refers to cells that putatively come from small clones (<75 cells); in the lower pie chart, grey refers to clonal labels that are represented in less than 1% of cells (Granulocytes). **C.** UMAP based on the static CpGs and the associated EPI-clone clusters for the old mouse. **D.** Comparison of clone sizes, measured using the percentage of cells in the clone in comparison to all cells, for the old/young mouse. Clones with less than 75 cells are shown in grey. **E.** Comparison of clone sizes for the 20 largest clones**. F.** Comparison of HSC/MPP1 output and myeloid output for the 20 clones with the highest HSC/myeloid output between the young and old mouse. **G**. Bubble plot visualizing the frequency (measured as the square of the HSC/MPP1 frequency) of HSC/MPP1 cells per clone for the old versus the young mouse. **H.** Visualizing clones in the differentiation uMAP for the old (left) and young (right) mouse. Two examples of clones are shown for both mice. **I, J.** Characterizing clonal behavior of all clones (rows) across the different cell types (columns) as a heatmap. The color indicates the relative frequency of the clones toward four main differentiation trajectories. Immature=HSC/MPP1, MPP2; myeloid=MPP4, myeloid progenitors; lymphoid=B cell progenitors; erythroid=MEP, erythroid progenitor. This is shown for the old (I) and young (J) mouse separately.

Decades of work have shown that aging provokes clonal expansion with profound loss of clonal diversity. However, much of our understanding of clonal behaviour in aged mammalian blood either comes from transplantation experiments^37^or mathematical modelling^38^, which may not recapitulate steady-state haematopoiesis or lacks the resolution of single-cell lineage analysis. More recently, transgenic barcoding mouse models have provided valuable insight into native hematopoietic development^10,39–41^, but these complex models have not been applied to ageing, due to experimental constraints and inaccessibility of the models. To tackle this question with EPI-clone, compared the data from the young mouse to a 100-week old mouse. Using the reference low-dimensional representation from the main experiment, we again identified the three differentiation trajectories, and weak shifts in cell type proportions between the young and the old mouse, including an increased ratio of HSCs to MPP3/4, which reflects known changes in the LSK compartment with age^42,43^ (Figure S5a, b). When comparing the EPI-clone output, we observed that in the old mouse 78.7% of cells were part of expanded clones, compared to 41.6% in the young mouse (Figure 4c,d, Figure S6, S7). Expanded clones in the old mouse were also larger, compared to the young mouse (Figure 4d, e). This gradual loss of clonality better mimics human aging, in stark contrast with results from barcoding transplantation experiments (cf. discussion)^44^.

Next, we measured the distribution of cell types for each clone across the various stem and progenitor clusters. Comparing the ratio of differentiating versus stem cells in the landscape, we first estimated the blood cell output of each of the stem cell clones. We observed a single, large, expanded clone containing mostly immature stem-like cells (Figure 4f, g, h). Five additional clones, smaller in size, also almost exclusively contained stem cells that appear to be dormant or incapable of proceeding with differentiation (Figure 4h, i, j). These clones resemble low-output stem cell clones previously observed in transplantation studies, particularly in old bone-marrow donors^11,45–49^. Next, we compared the various types of differentiating stem-cell progeny to quantify the changes in stem cell lineage biases. Old mice showed a very moderate increase in the number of myeloid-biased clones, in contrast with results from classic transplantation experiments (Figure 4f, i, j). However, the large, low-output stem cell clones were mostly myeloid biased. This result is in line with previous reports showing an age-dependent intrinsic myeloid shift and loss of differentiation potential on a per-stem-cell basis, and suggests that classic transplantation experiments mostly sampled stem cells from these highly self-renewing but rare low-output clones^47–49^. In summary, we show that EPI-clone can map high-resolution lineage trajectories in native hematopoiesis across the mouse lifespan and functionally characterize age-related changes in stem cell clone behaviors in a transgene-free manner.

### Various degrees of oligoclonality shape human blood ageing

Considering the success of EPI-clone to chart unperturbed mouse hematopoiesis, we decided to test its performance in human hematopoiesis. To characterize clonality in healthy human ageing, we designed a panel targeting 498 CpGs with variable methylation between or within blood progenitor populations^27,50^ (Figure S8a,b, Methods), 147 genomic regions commonly mutated in clonal hematopoiesis (CHIP), and 20 regions targeting chromosome Y. We then applied this panel to characterize total bone marrow from two male donors (71 and 77 years old). Samples were further stained with an antibody panel targeting 45 surface proteins to provide a phenotypic characterization. Analysis of the genotyping targets revealed that donor 1 had lost chromosome Y in a subset of cells, while donor 2 displayed no genotypic abnormalities (Figure 5a).

**Figure 5:**
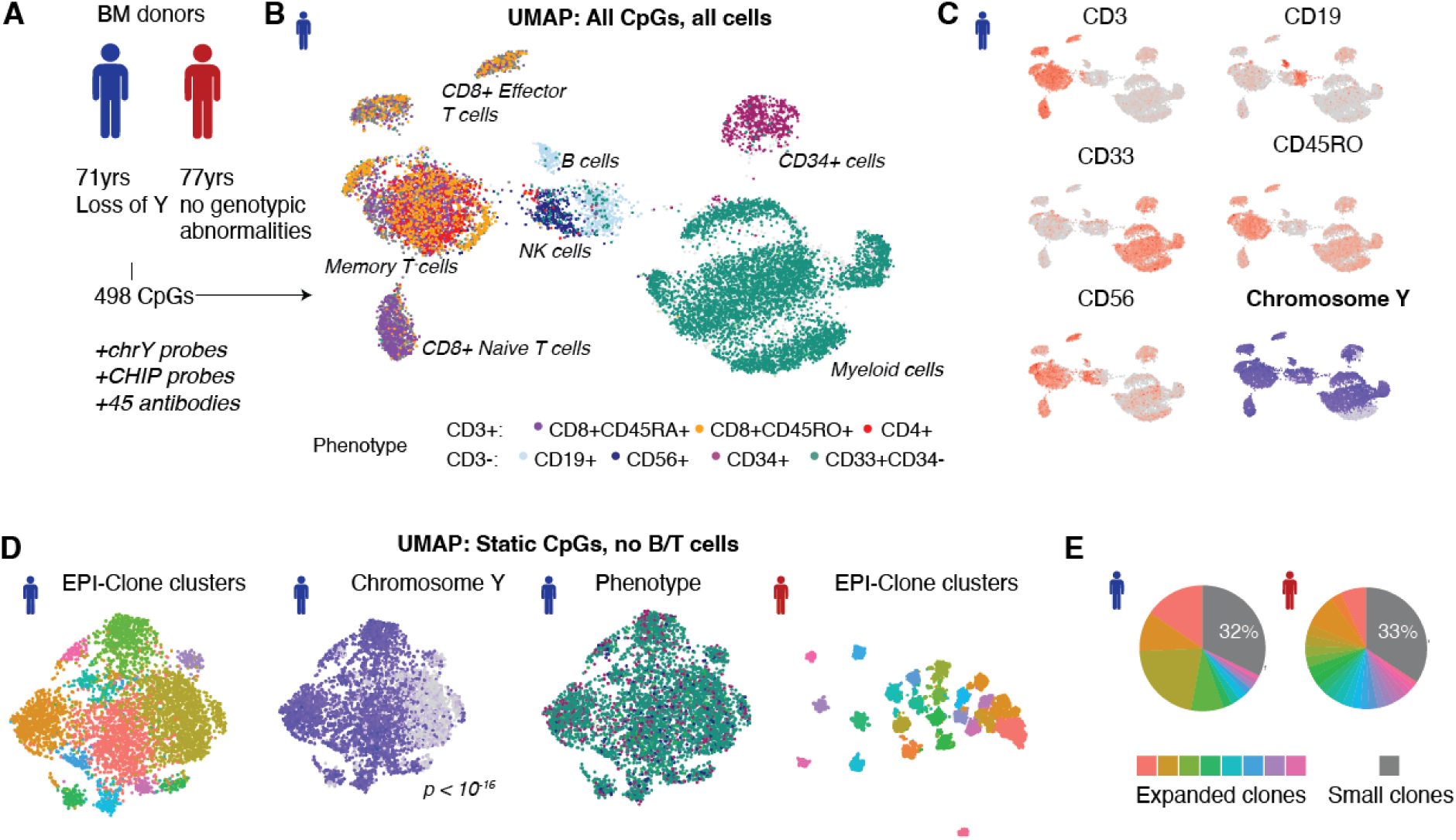
Different degrees of oligoclonality in human ageing. **A.** Experimental design. **B.** uMAP of DNA methylation data for donor 1. See Figure S8 for donor 2. **C.** Expression of surface proteins (red) and presence of chromosome Y amplicons (blue). **D.** EPI-Clone uMAP of static CpGs in all cells except B and T cells, see also Figure 3. From left to right: Donor 1, highlighting clone; Donor 1, highlighting loss of chromosome Y; Donor 2, highlighting clone. P-value is from a Fisher test for the enrichment of LoY in the expanded cluster. **E.** Fraction of total cells stemming from the different expanded clones.

As in mouse, both clonal identity (here defined by loss of chromosome Y [LoY]) and cell state (here defined by surface phenotype) jointly influenced the global methylation profile (Figure 5b, c, Figure S8c). LoY occurred in parts of the myeloid cells, CD34+ cells and NK cells. Within each of these cellular compartments, LoY cells clustered apart from the remaining cells. To further separate clonal identity from cell state information present in the DNA methylation data, we applied the EPI-Clone algorithm (Figure S8d,e). In the case of donor 1, EPI-Clone identified 11 clones that, individually, contributed to up to 20% of blood formation. All cells within the largest clone showed LoY, while LoY was also observed in 3 further (sub-)clones, suggesting independent acquisition of these lesions (Figure 5d, e). These results mimic recent reports using colony-based whole genome sequencing in 70-80 year-old human hematopoietic cells, but at a small fraction of the cost and with simultaneous information on cell states^13^. In the case of donor 2, EPI-Clone identified 23 expanded clones that individually contributed to 1.3-6.9% of blood formation, indicating significantly less oligoclonal blood production (Figure 5d, e). Together, these results demonstrate the capability of EPI-Clone to identify hematopoietic clones in humans, and they demonstrate variable, oligoclonal blood formation in individuals older than 70 years.

## Discussion

Here, we have demonstrated that DNA methylation at a few hundred CpGs is sufficient to simultaneously identify clones and provide detailed information on the cellular state at the level of individual hematopoietic cells. Targeted scDNAm profiling with single-cell, single-CpG resolution permits to capture both layers of information in a high-throughput commercial assay, enabling studies of clonal dynamics and function without need for genetically-engineered mouse models. Ground truth clonal labels are well recapitulated in both mouse experiments (using genetic lineage tracing barcodes as ground truth) and in the human study (using loss of chromosome Y as ground truth). This indicates that clonal somatic epimutations are stable over extended periods of time. In mouse, four to six months elapsed between induction of the ground truth clonal label and the harvesting of cells post-transplantation. In human, previous studies indicate that decades pass between the initial loss of Y and the observation of expanded LoY clones in age^13^. Clonal information was further stable through multiple cell divisions from the stem cell until terminal differentiation. Together, somatic epimutations appear to be a uniquely stable, long-term lineage tracer.

The robustness of EPI-clone is best evidenced by its capacity to identify high-resolution clonal patterns in native murine hematopoiesis. Using EPI-clone, here we provided a first picture of *in situ* high-resolution clonal dynamics in mouse aging. Mimicking the clonal evolution in human hematopoiesis, murine hematopoiesis shifts from highly polyclonal to oligoclonal blood production, although retaining more overall diversity than in humans^13^. While aged HSCs have been traditionally characterized by functional loss of lymphoid potential and inefficient differentiation, we find that *in situ* these behaviors are notably restricted to a small number of HSC clones, with the majority of clones behaving just like in the young age^51^. These rare low-output clones make up a large fraction of the total cells of the HSC compartment (∼30%), suggesting that these cells are a major contributor to the phenotypes observed in transplantation studies. Thus, our observations are compatible with the exhausted stem cell theory, which postulates that HSCs functionally decline with subsequent activations and cell divisions, but support that aging is highly heterogeneous at the clonal level, with many clones protected from age-related decline^52^. In the future, EPI-clone may provide clues to the mechanisms of inflammation-driven age acceleration as well as become a platform to test immune rejuvenation interventions^53^. Critically, we also provide a proof of principle that EPI-clone enables the low-cost and high-throughput characterization of hematopoietic stem cell clones during aging in humans, where genetic barcoding is not applicable, allowing future studies of samples from clinical trials or epidemiological studies.

A final important point relates to where and how epimutations arise and label HSC clones, becoming stable tracers for clonal behaviors. We found that they (i) randomly occur, but remain stable over many cell divisions, (ii) are enriched for heterochromatic and late-replicating domains, and (iii) mainly arise from a fully methylated default state. As a potential explanation, in fast-dividing cells DNMT1 may act insufficiently to copy the DNA methylation state to the nascent DNA strand, especially in late-replicating domains. Consequently, if the cell divides before the copying of DNA methylation, one of the daughter cells will lose the DNA methylation state of the cell of origin (Figure S9). This epimutation will then be passed to all progeny of the cell, especially if it occurs in a silent heterochromatic region without direct functional consequence. A preprinted study of bulk methylome profiles from blood cells in monozygotic twins suggests that clone-associated variation of the methylome may be established during embryonic development^25^. This would be consistent with the long-term stability of the clonal mark observed in our study.

### Limitations

While we found that both clonal identity and differentiation state can be identified using only 453 CpGs, the targeted nature of scTAM-seq requires the availability of bulk methylome resources and can lead to lower cell-type resolution of some cell types, as observed here in the context of the mouse late myeloid precursors. However, cell-types can be further differentiated based on surface markers. A further limitation of EPI-Clone is that only cells belonging to expanded clones can be assigned to their clone of origin. Cells belonging to clones contributing less than 0.2-1% of cells are identified as not belonging to expanded clones, but their clonal identity can currently not be inferred.

## Data Availability

The scDNAm dataset (https://doi.org/10.6084/m9.figshare.24204750) and the scRNA-seq dataset (https://doi.org/10.6084/m9.figshare.24260743.v1) are available as Seurat objects from Figshare.

## Code Availability

Code for processing scTAM-seq data, for the EPI-clone algorithm, and for generating all figures of the paper is available from GitHub (https://github.com/veltenlab/EPI-clone).

## Acknowledgements

The authors would like to thank Renée Beekman, Roser Zaurin and Agostina Bianchi for collaboration on the scTAM-seq method, Mission Bio for support with panel designs and the CRG Genomics Unit for assistance with sequencing. This work was funded by the European Hematology Association (EHA Advanced Research Grant to L.V.) and the Asociación Española Contra el Cáncer (AECC lab grant to L.V.). M.S. was supported through the Walter Benjamin Fellowship funded by Deutsche Forschungsgemeinschaft (DFG, German Research Foundation, reference 493935791). I.S. was supported through the European Union’s Horizon 2020 research and innovation programme under the Marie Skłodowska-Curie grant agreement No 945352. L.V. acknowledges support of the Spanish Ministry of Science and Innovation to the EMBL partnership, the Centro de Excelencia Severo Ochoa and the CERCA Programme / Generalitat de Catalunya. A.R.F has been supported by the Cris Foundation Excellence Award (PR_EX_2020-24), the ERC Starting Grant MemOriStem (101042992), the Spanish National Research Agency (PID2020-114638RA-I00), the Agencia de Gestio d’Ajuts Universitaris i de Recerca (AGAUR, 2017 SGR 1322), and the CERCA Program/Generalitat de Catalunya. A.R.F. acknowledges support from the Institut Catalá de Recerca i Estudis Avançats (ICREA), the American Society of Hematology (ASH) Scholar Award, the Leukemia Lymphoma Society Special Fellow Career Development Program Award (3391–19), the NIH NHLBI K99/R00 transition to independence award (K99 HL146983), the Ministry of Science Ramon y Cajal Fellowship, and the LaCaixa Junior Fellows Incoming Fellowship.

## Author contributions

I.S. performed the mouse experiments. C.S.T. and I.S. generated all sequencing libraries. M.S., M.B., and L.V. analyzed the data and developed EPI-Clone, with conceptual input from I.S. and A.R.F.. M.S. and M.B. created the targeting panels with support by R.F.. J.R. and S.B.C analyzed the scRNA-seq data. M.K. and S.R. provided and characterized the human samples. M.S., I.S., A.R.F, and L.V. conceptualized the study. M.S., I.S., A.R.F., and L.V. wrote the manuscript with input from all co-authors.

## Online Methods

### Mice and Animal Guidelines

All procedures involving animals adhered to the pertinent regulations and guidelines. Approval and oversight for all protocols and strains of mice were granted by the Institutional Review Board and the Institutional Animal Care and Use Committee at Parque Cientifico de Barcelona under protocols CEEA-PCB-22-001-ARF-P1; CEEA-PCB-22-002-ARF-P2. The study is following all relevant ethical regulations. CD45.1 (CD45.1, B6.SJL-Ptprca Pep3b/BoyJ, stock no. 002014, The Jackson Laboratory) mice were used as transplantation recipients for CD45.2 (BL6/J) donor cells. Both male and female mice, 6-12 weeks of age, were used as donors and recipients. Mice were kept under specific-pathogen free conditions for all experiments.

### Hematopoietic stem cell isolation

Following euthanasia, bone marrow was harvested from the femur, tibia, pelvis, and sternum through mechanical crushing, ensuring the retrieval of most of the cells. The collected bone marrow cells were then sieved through a 40-μm strainer and cleansed with a cold ’Easy Sep’ buffer containing PBS, 2% fetal bovine serum (FBS), 1 mM EDTA, and Penicillin/Streptomycin followed by lysis of red blood cells using RBC lysis buffer (Biolegend, Catalog no. 420302). At first, mature lineage cells were selectively depleted through the Lineage Cell Depletion Kit, mouse (Miltenyi Biotec, Catalog no. 130-110-470), while the resulting Lin^-^ (lineage-negative) fraction was then enriched for c-Kit expression using CD117 MicroBeads (Miltenyi Biotec, Catalog no: 130-091-224). These cKit-enriched cells were washed, blocked with FcX and stained with following fluorescently labeled antibodies: APC anti-mouse CD117, clone ACK2 (Biolegend catalog no. 105812), PE/Cy7 anti-mouse Ly6a (Sca-1) (Biolegend, catalog no. 108114); Pacific Blue anti-mouse Lineage Cocktail (Biolegend, catalog no. 133310); PE anti-mouse CD201 (EPCR) (Biolegend, catalog no. 141504); PE/Cy5 anti-mouse CD150 (SLAM) (Biolegend, catalog no. 115912); APC/Cyanine7 anti-mouse CD48 (Biolegend, catalog no. 103432). For transplants, EPCR^+^Lin^-^Sca-1^+^c-Kit^+^ HSCs were sorted via fluorescence-activated cell sorting (FACS) employing a BD FACSAria Fusion with a 70uM nozzle. For single-cell analyses, the c-Kit enriched fraction of cells was additionally stained with the TotalSeq-B antibody cocktail (Table S1), and different compartments were sorted as outlined in Figure S1c.

### Mature immune cell experiment

For this study, the mouse was anesthetized and perfused. Post perfusion the lungs were extracted from the chest cavity, and a single-cell suspension was prepared through protease and DNAse solution in the Lung Dissociation Kit (Miltenyi Biotech, catalog #130–095-927) followed by mechanical dissociation using gentleMACS “C” columns (Miltenyi Biotech, catalog #130–093-237) according to the manufacturer’s instructions. The dissociated cells were filtered using a 70 μm strainer and centrifuged at 400 g for 5 min at room temperature. The supernatant was removed by aspiration and red blood cell lysis was performed using RBC lysis buffer (Biolegend, Catalog no. 420302). Cells were then washed with FACS buffer and pelleted at 400 g for 5 min at 4°C. The supernatant was removed, and the pellet was resuspended in Fluorescence-Activated Cell Sorting (FACS) buffer before being passed through a 40 μm strainer and stained for the mature myeloid cell marker. Sorted cells were then stained for the TotalSeq-B antibody cocktail together with LSK, LKs, and mature cells from bone marrow and loaded into the Mission Bio Tapestri platform at the recommended concentration.

### Old and Young native hematopoiesis experiment

For this study, LSK and LKs were extracted and stained as mentioned earlier for mission bio run from young 12-week-old BL6/J (CD45.2) mice and old 100-week-old BL6/J (CD45.2) mice.

### Transplantation Assay

HSCs were harvested from 8-week-old BL6/J (CD45.2) mice and transduced with GFP or T-Sapphire-tagged LARRY barcode lentiviral libraries and subsequently transplanted into CD45.1 recipient mouse via tail-vein injection with 150,000 whole bone marrow cells as support cells, with each recipient mouse receiving a 150 μl volume of PBS injection. The CD45.1 recipient mouse had been preconditioned with a lethal X-ray radiation dose, administered as two separate sessions of 5 Gy each, with a 2-hour interval between them. To assess the engraftment of donor cells, the percentage of CD45.2+ peripheral blood leukocytes (and the percentage of fluorescent-protein-labeled cells) in the recipient mice was determined. All mice demonstrated stable long-term engraftment until the experimental endpoint. Engraftment analysis, along with the measurement of labeling frequency, was carried out using BD FACS Fusion.

In vitro cultures of HSCs were done under self-renewing F12-PVA based conditions as described previously^54^. To prepare for cell sorting, 96-well flat-bottom plates from Thermo Scientific were coated with a layer of 100 ng/ml fibronectin (Bovine Fibronectin Protein, CF Catalog: 1030-FN) for 30 minutes at room temperature. Following the sorting process, HSCs were transferred into 200 µl of complete HSC media supplemented with 100ng/ml recombinant mouse TPO and 10ng/ml recombinant mouse SCF (PeproTech Recombinant Murine TPO Catalogue Number: 315-14; PeproTech Recombinant Murine SCF, Catalogue Number: 250-03) and grown at 37°C with 5% CO2. During lentiviral library transduction, the first media change took place 24 hours post-transduction. Three days post-labeling, the cultured HSCs were collected and subsequently transplanted into CD45.1 mice.

### Construction of lentiviral pLARRY vectors

The construction of barcoded libraries was executed in accordance with a previously established protocol (https://www.protocols.io/view/barcode-plasmid-library-cloning-4hggt3w). First, the T-Sapphire or EGFP coding sequences, and the EF1a promoter sequences were PCR amplified from pEB1-T-Sapphire and pLARRY-EGFP with primers homologous to the vector insertion site in a custom lentiviral plasmid backbone (Vectorbuilder, Inc) using Gibson assembly (Gibson Assembly® Master Mix, NEB, Ref. E2611L). After magnetic-bead purification, ligated vectors were then transformed into NEB10-beta electroporation ultracompetent *E.coli* cells (NEB® 10-beta Electrocompetent E. coli, NEB, Ref.C3020K) and grown overnight on LB plates supplemented with 50 μg/mL Carbenicillin (Carbenicillin disodium salt, Thermo Scientific Chemicals Ref. 11568616). Colonies were scrapped using LB medium and pelleted by centrifugation. Plasmid maxipreps were performed using the Endotoxin-Free Plasmid Maxi Kit (Macheray Nagel), following the manufacturer protocol. pEB1-T-Sapphire was a gift from Philippe Cluzel (Addgene plasmid 103977). pLARRY-EGFP was a gift from Fernando Camargo (Addgene plasmid 140025)

### Barcode lentivirus library generation and diversity estimation

To barcode pLARRY plasmids and generate a library, a spacer sequence flanked by EcoRV restriction sites was cloned into the plasmid after the WPRE element of the vector. Custom PAGE-purified single-strand oligonucleotides with a pattern of 20 random-bases (GTTCCANNNNTGNNNNCANNNNGTNNNNAGNNNN) and surrounded by 25 nucleotides homologous to the vector insertion site were synthesized by IDT DNA Technologies. The assembly of these components and subsequent purification steps were carried out through Gibson assembly (Gibson Assembly® Master Mix, NEB, Ref. E2611L). Six electroporations of the bead-purified ligations were performed into NEB10-beta *E.coli* cells (NEB® 10-beta Electrocompetent E. coli, New England BiolabsEB, Ref.C3020K) utilizing a Gene Pulser electroporator (Biorad). Subsequently after transformation the cells were incubated at 37 degrees for 1 hour at 220 rpm. Post-incubation, the transformed cells were plated in six large LB-ampicillin agar plates overnight at 30°C. Colonies from all six plates were collected by scraping with LB-ampicillin and then grown for an additional 2h at 225 rpm and 30 °C. Cultures were pelleted by centrifugation, and plasmids were isolated using the Endotoxin-Free Plasmid Maxi Kit (Macheray-Nagel), following the manufacturer protocol.

For estimating diversity, barcode amplicon libraries were prepared by PCR amplification of the lentiviral library maxiprep using flanking oligonucleotides carrying TruSeq read1 and read2 adaptors using 10 ng of the library (Table S2). We used the minimal number of cycles that we could detect by qPCR in order to avoid PCR amplification bias (10-12 cycles). After bead purification, 10 ng of the first PCR product was used as a template for a second PCR to add Illumina P5 and P7 adaptors and indexes (Table S2). Two independent PCRs were sequenced on an Illumina NovaSeq 6000 S4 platform (Novogene UK) to confirm diversity after correction of errors through collapsing with a Hamming-distance of 4. After collapsing, libraries were confirmed to contain at least 50 million different barcodes, with enough diversity for uniquely labeling up to 100,000 HSCs with a minimal false-positive rate.

### Lentivirus production and Barcode labeling

Lentivirus production and HSPC transduction was performed as described Weinreb et al.^2^

### Human samples

Bone marrow samples from healthy volunteers were obtained at the Heidelberg University Hospital after informed written consent using ethic application number S-480/2011. All experiments involving human samples were approved by the ethics committee of the University Hospital Heidelberg and were in accordance with the Declaration of Helsinki.

Samples from healthy donors were thawn and stained using CD34 sorting antibody (BioLegend, 343517) and a pool of oligo-conjugated antibodies from the TotalSeq-D Heme Oncology Cocktail from BioLegend (MB53-0053) as well as two additional TotalSeq-D antibodies from BioLegend (CD135, 313325; CD49f, 313641). Samples were then sorted for CD34+ and CD34-populations subjected to scTAM-seq. For Donor 1, 102,000 CD34+ cells and 298,000 CD34-cells were pooled and, from this pool, 78,050 cells were loaded onto the Tapestri instrument. For Donor 2, 91,006 CD34+ cells and 476,358 CD34-cells were pooled and, from this pool, 90,650 cells were loaded onto the instrument.

### Single-cell DNA methylation profiling and single-cell RNA-seq

For profiling DNA methylation at single-cell resolution, we used scTAM-seq^30^, which leverages the Mission Bio Tapestri technology to investigate up to 1,000 CpGs in up to 10,000 cells per experiment. Briefly, we loaded 120,000-140,000 cells into the Tapestri machine and followed the default Mission Bio DNA+Protein protocol for V2 chemistry (https://missionbio.com/wp-content/uploads/2021/02/Tapestri-Single-Cell-DNA-Protein-Sequencing-V2-User-Guide-PN_3360A.pdf), with modifications: (i) we added a DNA methylation sensitive restriction enzyme (HhaI) to remove non-methylated targets prior to amplification. (ii) we used TotalSeq-B antibodies and different primers for the amplification of antibody oligonucleotide tags. The default Mission Bio protocol uses a different type of oligonucleotide tag, TotalSeq-D, which are currently not available for mouse antigens. The aging experiment was performed with Tapestri workflow v3 (https://missionbio.com/wp-content/uploads/2023/08/Tapestri-Single-Cell-DNA-Protein-Sequencing-v3-User-Guide_MB05-0018.pdf).

In detail, we added 5μL of highly-concentrated HhaI (150,000U/mL, NEB) enzyme and 5 µL of 30 µM of a custom Antibody Tag Primer specific for the amplification of the oligonucleotide tags of TotalSeq-B antibodies (ACTCGCAGTAGTCTTGCTAGGACCGGCCTTAAAG) to the Tapestri Barcoding Mix V2 reagent. An incubation at 37°C for 30 minutes was added to the start of the Targeted PCR thermal cycling program to allow for the restriction enzyme digest to take place prior to the PCR amplification step. The use of TotalSeq-B antibodies primarily affected the “Protein Library Cleanup I” section of the protocol where we replaced the 2X Binding & Washing (B&W) Buffer from the kit with the following buffer prepared with nuclease-free water: Tris-HCl (final concentration 10 mM, pH7.5), EDTA (final concentration 1 mM), and NaCl (final concentration 2 M). We used 2 µL of 5 µM of our custom Biotin Oligo (/5Biosg/GTGACTGGAGTTCAGACGTGTG/3C6/) to isolate the antibody tags. In addition, during the isolation of antibody tags, we performed the second wash of Streptavidin Beads with 1 mL nuclease-free water instead of 1X B&W Buffer. Finally, each tube of Steptavidin Beads was resuspended in 45 µl of nuclease-free water then transferred and combined into a new tube for a total of 90 µl. To amplify the final protein target library, we used 5 µL of 4 µM of each custom indexed primers (Forward: CAAGCAGAAGACGGCATACGAGAT[i7 index]GTGACTGGAGTTCAGACGTGTGCTCTTCCGATCT 3’, Reverse: AATGATACGGCGACCACCGAGATCTACAC[i5 index]TCGTCGGCAGCGTC). Typically, we performed twice as many reactions to amplify the DNA target library, but this may be increased to achieve sufficient yield. Lastly, we adjusted the AMPure XP reagent to sample ratio in the second size selection step in “DNA Library Cleanup II” from 0.72X to 0.65X.

Using the stained cells that we used as input to scTAM-seq (Figure S1c), we additionally performed 10x Genomics Chromium Single Cell 3’ for transcriptomic profiling of the cells, following the standard protocol. LARRY barcodes were later amplified using a modified version of the protocol described in Weinreb et al. ^2^ (see Table S2 for an updated list of primers).

### Mouse panel design for scTAM-seq

We aimed at designing a panel harboring CpGs dynamically methylated in HSCs, as well as in more committed progenitors (MPPs). We collected bulk whole-genome bisulfite sequencing data from a previous publication^28^ profiling DNA methylation in three replicates of HSCs (LSK=Lin^-^Sca-1^+^c-KIT^+^ and CD135^-^CD48^-^CD150^+^CD34^-^), MPP1 (LSK, CD135^-^CD48^-^CD150^+^CD34^+^), MPP2 (LSK, CD135^-^CD48^+^CD150^+^CD34^+^) and a mixture of MPP3 (LSK, CD135^-^CD150^-^CD48^+^CD34^+^) and MPP4 (LSK, CD135^+^CD150^-^CD48^+^CD34^+^). Using this data, we selected CpGs from three classes that are variably methylated in HSPCs (Figure S1c,d): (i) CpGs differentially methylated between the HSCs and the different MPP populations, (ii) CpGs intermediately methylated within HSCs, and (iii) CpGs harboring within-sample heterogeneity in HSCs. The code for selecting CpGs is available from https://github.com/veltenlab/EPI-CloneSelection

i. *Differentially methylated CpGs (DMCs).* We used RnBeads^31^ to determine CpGs that are specifically methylated in one of the HSPCs (i.e., in either HSC, MPP1, MPP2, MPP3/4), while being unmethylated in all the remaining HSPC populations. We only focused on CpGs that were covered by at least 10 sequencing reads in all samples and that had a methylation difference of at least 0.2 between the target cell type and the average of the remaining cell types. After sorting the CpGs by the methylation difference between the cell types, we investigated whether the CpG is located in the HhaI cut-sequence (GCGC) and annotated for vicinity to important hematopoietic TFBS.
ii. *Intermediately methylated CpGs (IMCs).* IMCs have to be non-overlapping with DMCs and are then defined by a DNA methylation level in the bulk samples between 0.25 and 0.75. Such CpGs may be differentially methylated between two sub-cell types of HSCs. IMCs are required to have a low Proportion of Discordant Reads (PDR^55^) together with a high quantitative Fraction of Discordant Read Pairs (qFDRP^32^). PDR and qFDRP are measures of within-sample heterogeneity (WSH) in bulk sequencing data and quantify the concordance of methylation states on the same sequencing read (PDR) or of multiple CpGs across different sequencing reads (qFDRP).
iii. *CpGs harboring within-sample heterogeneity (WSH).* CpGs with high WSH are non-overlapping with DMCs and IMCs. The CpGs are then identified based on high levels of both PDR and qFDRP. These CpGs thus are located in regions showing variable methylation profiles in bulk sequencing data and might represent regions with stochastic methylation in HSCs.

After identifying all CpGs fulfilling the criteria above, we enriched the selected CpGs for those located in the vicinity (100bp) of at least one TFBS of an important hematopoietic TF (Table S3). We then selected 105 CpGs specifically methylated in HSCs, 70 in MPP1, 70 in MPP2, 75 in MPP3/4, 210 IMCs, and 80 WSH (Figure S1c,d). Additionally, we included the following control amplicons: 20 constitutively methylated, 20 constitutively unmethylated and 50 amplicons without a HhaI cut-sequence. Control amplicons are required to identify cells from the data, because the remaining amplicons are digested depending on their methylation state. We uploaded this list to the Mission Bio Designer tool (https://designer.missionbio.com/) to receive a final list of 663 amplicons and corresponding primer sequences (Table S3). The CpGs were further annotated according to their location in the genome with respect to chromatin states defined in Vu et al., 2023^33^.

### Human panel design for scTAM-seq

The design for the human panel closely followed the guidelines of the mouse panel. Two previously published datasets were used to similarly profile DMCs, IMCs, and WSH. Sites were selected to not include single nucleotide polymorphisms according to dbSNP version 151 and to be located in the HhaI cut sequence.

i. *Differentially methylated CpGs (DMCs).* We considered peripheral blood and bone marrow samples from the dataset of Farlik et al^27^. Samples with an average coverage across all CpGs below 1 were removed. Differentially methylated CpGs between HSCs, MPPs, multi-lymphoid progenitors (MLPs; combining MLP0, MLP1, MLP2, MLP3), common lymphoid progenitors (CLPs), common myeloid progenitors (CMPs), granulocyte-macrophage progenitors (GMPs) were computed using *RnBeads*. CpGs with a mean methylation difference higher than 0.1 between the cell types were identified as DMCs.
ii. *Intermediately methylated CpGs (IMCs).* We performed IMC detection on HSC-enriched lineage-negative (Lin− CD34+ CD38−) samples from eight male donors using the Adelman et al.^50^ dataset. To deal with data sparsity, we set the maximum quantile of missing values per site to 0.005 and removed any sites that exceeded this threshold. IMCs were defined as CpGs with a DNA methylation level between 0.25 and 0.75 in at least 5 samples. When checking for a HhaI cut site we allowed for a maximum of 25 CpG sites in the extended region around the IMC.
iii. *CpGs harboring within-sample heterogeneity (WSH).* We used the same data set to identify CpGs with a variance higher than 0.1 across all samples.

We additionally created genotyping amplicons that cover mutations in ASXL1, DNMT3A, TET2, TP53, JAK2, IDH2, PPM1D, SF3B1, IDH1, and SRSF2. We used 62 amplicons covering these genes from the Tapestri Single-cell DNA Myeloid Panel by Mission Bio (https://missionbio.com/products/panels/myeloid/) as a base panel, excluding amplicons harboring a the HhaI restriction sequence *GCGC*. We designed further amplicons for exons in the aforementioned genes that had a coverage of less than 60% in the default myeloid panel. To prevent these amplicons from harboring a recognition site, we perfomed a virtual digestion of the exonic sequences using the HhaI cut sequence. We then uploaded a list containing the fragmented genomic regions to the Mission Bio Designer tool, which resulted in 82 additional amplicons. We included 20 amplicons targeting chromosome Y designed according to the additional genotyping amplicons and 50 control amplicons without an HhaI cut-sequence. We uploaded the CpG targets and readily designed genotyping, chromosome Y, and control amplicons using the Mission Bio Designer tool. The final list comprises 665 amplicons and corresponding primer sequences. The resulting 448 CpG targeting amplicons are divided into 215 DMC, 145 IML, and 88 WSH amplicons (Table S4).

### Sequencing

Libraries were sequenced on an IIllumina NovaSeq 6000 with 2×100bp (scTAM-seq mouse), 2×150bp (scTAM-seq human), 2×50 bp (scRNA-seq) and 2×50 bp (protein libraries) reads. For an overview of the sequencing statistics see Table S5.

### Data processing

For processing of raw data we used the pipeline available at https://github.com/veltenlab/scTAM-seq-scripts. Briefly, barcodes were extracted from the raw sequencing files before alignment to the reference genome subset to the CpG panel. Reads mapping to each of the amplicons were quantified to generate a count matrix and DNA methylation states were determined using a cutoff of one sequencing read as in the original scTAM-seq publication^30^. We used those cellular barcodes that had more than 10 sequencing reads in at least 70% of the control (non-HhaI) amplicons. Doublets were removed using the DoubletDetection tool (https://zenodo.org/record/2678042).

To determine the primer combinations that reliably amplify in our panel, we performed a single experiment without the restriction enzyme. For this experiment, wildtype Lin^-^cKIT^+^ cells were used and we determined that 453 of the 573 non-control amplicons (79%) amplified in more than 90% of the cells. These amplicons were used for subsequent analysis.

For the surface protein data, the Mission Bio pipeline was used to extract sequencing reads for a particular cell-barcode/antibody-barcode combination. We restricted analysis of the protein data to those cellular barcodes identified in the DNA methylation library.

### Data integration and annotation of cell states

We constructed Seurat^56^ objects for each of the scTAM-seq samples individually using the binary DNA methylation matrix. To integrate all the samples in the main LARRY experiment, we used Seurat’s IntegrateData^56^ function. Then, we used Seurat’s standard workflow without normalization to obtain a low dimensional representation of our data using uMAP. We removed cells in low-density parts of the uMAP, since we found these cells to be of lower quality using the non-digested control amplicons. To annotate the cell type clusters we obtained as result of the Seurat workflow, we inspected (i) the expression of surface protein, (iii) the DNA methylation states of important lineage-specific TFs, and (iii) the DNA methylation states of CpGs in bulk data. To that end, we first determined sites unmethylated in a cell type cluster using the Wilcoxon Rank Sum test. For those sites, we investigated whether they are in the vicinity of any of the 39 TFs in Table S3 and computed enrichment p-values with the Fisher exact test. A full vignette is available at https://github.com/veltenlab/EPI-clone.

In the case of the mature cell and aging experiments, we did not perform batch integration, since both samples were processed as one batch. To remove the effect of clone on the differentiation UMAP, we computed the low-dimensional representation only on CpGs not significantly (Chi-Square test p-value after Bonferroni correction larger than 1×10^-8^) associated with the LARRY barcode. Annotation of the cell type clusters was analogously performed using bulk methylation values, de-methylation of TFBS, and surface protein expression.

For the aging experiment, we used the static CpGs selected in the main LARRY experiment to perform clonal clustering. We then selected the clones with more than 75 cells, due to the larger overall cell number in the aging experiment (Mission Bio Tapestri version v2 vs. v3, Table S5). For comparing the clonal output across the two experiments, we quantified the contribution of each clone towards the three lineages and visualized this as a heatmap or as barcharts.

For the human experiment, the data of each patient was investigated separately. Analysis was conducted as described above, but cell types were annotated exclusively using surface protein expression.

For the scRNA-seq dataset, we used cellranger to generate transcriptomic and surface protein count matrices, which were used as input to Seurat. Harmony^57^ was used for batch integration and the cell type annotation was performed using known hematopoietic marker genes together with the expression of surface proteins.

### Processing of LARRY Barcodes

Sequencing reads mapping to the amplicon harboring the LARRY barcode were extracted from the raw sequencing reads using the fluorophore sequence:

GCTAGGAGAGACCATATGGGATCCGAT. The LARRY barcode was determined using the base pairs following the GFP sequence, given that the sequence matches the rules by which the barcode was constructed. Barcode extraction was performed using a modified version of the scripts provided in the original LARRY publication^2^ (https://github.com/AllonKleinLab/LARRY). Barcodes supported by less than 5 sequencing reads were discarded and LARRY barcodes with a Hamming distance lower than 3 were merged for each of the experimental batches individually.

Notably, each cell can harbor more than one unique LARRY barcode due to multiple lentiviral infections. In these cases, groups of LARRY barcodes get jointly passed on to the progeny. To call clones in this setting, we computed for any pair of LARRY barcodes the extent to which these two barcodes were observed in an overlapping set of cells (formally, a Jaccard index). LARRY barcodes were then clustered according to this distance metric. We used a permutation test to determine LARRY barcodes that are merged together to a clone. When LARRY barcodes were merged, cells were assigned to the merged clone if any constituent LARRY barcode was observed.

### Clustering by clone (EPI-Clone)

The EPI-Clone algorithm is divided into three steps, (i) identification of static CpGs, (ii) identification of cells from expanded clones and (iii) clustering of cells from expanded clones. A detailed, step-by-step vignette is available at https://github.com/veltenlab/EPI-clone. In brief:

i. *Identification of static CpGs*. For each combination of CpG and surface protein, EPI-Clone performs a Kolmogorov-Smirnov test to investigate if cells with methylated CpG differ in surface antigen expression relative to cells with unmethylated CpG. CpGs with no significant antigen association according to a Bonferroni criterion are then selected, if their average methylation across all cells is less than 90% but higher than 25% in mouse and higher than 5% in human. In the main LARRY experiment from figure 2/3, this resulted in the identification of 110 CpGs. In the human experiment depicted in figure 5, 113 CpGs were identified.
ii. *Identification of cells from expanded clones.* PCA is performed on all CpGs from step (1). In the reduced dimensional space obtained by the first n=100 PCs, the average Euclidean distance to the k=5 nearest neighbors is determined. Effects of cell state, batch and sequencing depth on this measure of local density are then removed by linear regression. We observed that smoothing the resulting quantity locally over twenty nearest neighbors additionally improved performance. Optimal parameters n and k of this step were identified by systematic grid search, using LARRY barcodes as a ground truth; they can also be justified by the consideration that n needs to be large (since there is little co-variation between stochastic clonal epi-mutations), and k needs to be small (since the goal here is to identify cells interspersed between clusters of clones).
iii. *Clustering of expanded clones.* Cells from expanded clones were clustered using the standard Seurat workflow, again in a space spanned by n=100 PCs.

## Supplementary Figures

**Figure S1:**
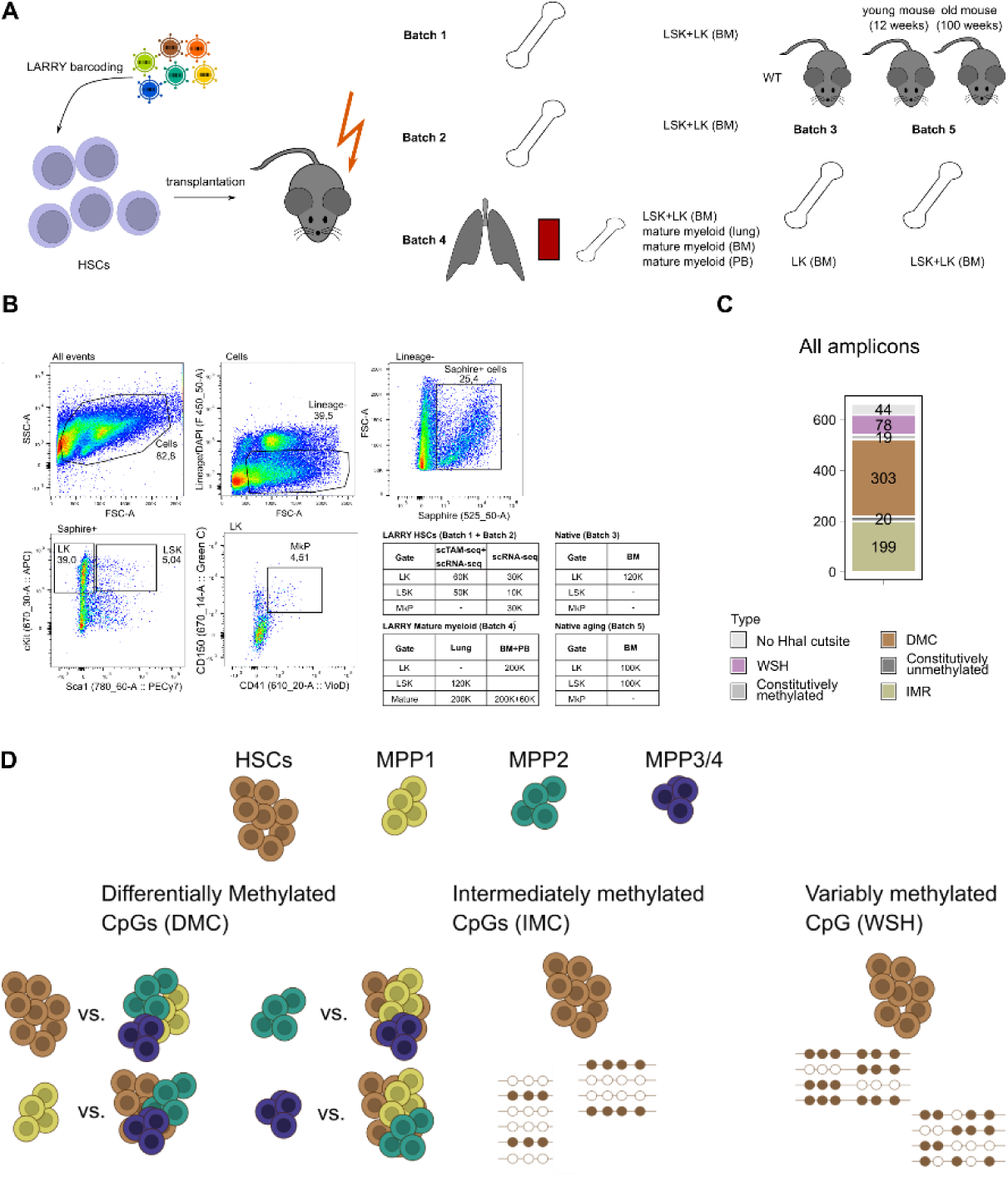
Overview of experimental design, CpG panel and FACS scheme. **A.** Overview of experimental design. Data generation involved five experimental batches (i) main LARRY experiment (LK, LSKs), (ii) replicate LARRY experiment (LK, LSK), (iii) native hematopoiesis (LK), (iv) mature myeloid cells from lung, bone marrow (BM) and peripheral blood (PB), and (v) aging experiment for native hematopoiesis. **B.** FACS scheme employed for the main LARRY experiments. The table shows the number of cells from different gates used as input for the experiments. **C.** Distribution of the CpGs covered by all 663 amplicons in our panel. For abbreviations, see panel D. **D.** Schematic overview of the CpG selection for scTAM-seq. Bulk DNA methylation data was collected from Cabezas-Wallscheid et al., 2014. We identified three classes of CpGs, which we included in the final panel design shown in Figure 1b: DMCs, IMCs, and WSH. The lines represent sequencing reads, where filled circles stand for methylated and unfilled circles for unmethylated CpGs, respectively.

**Figure S2:**
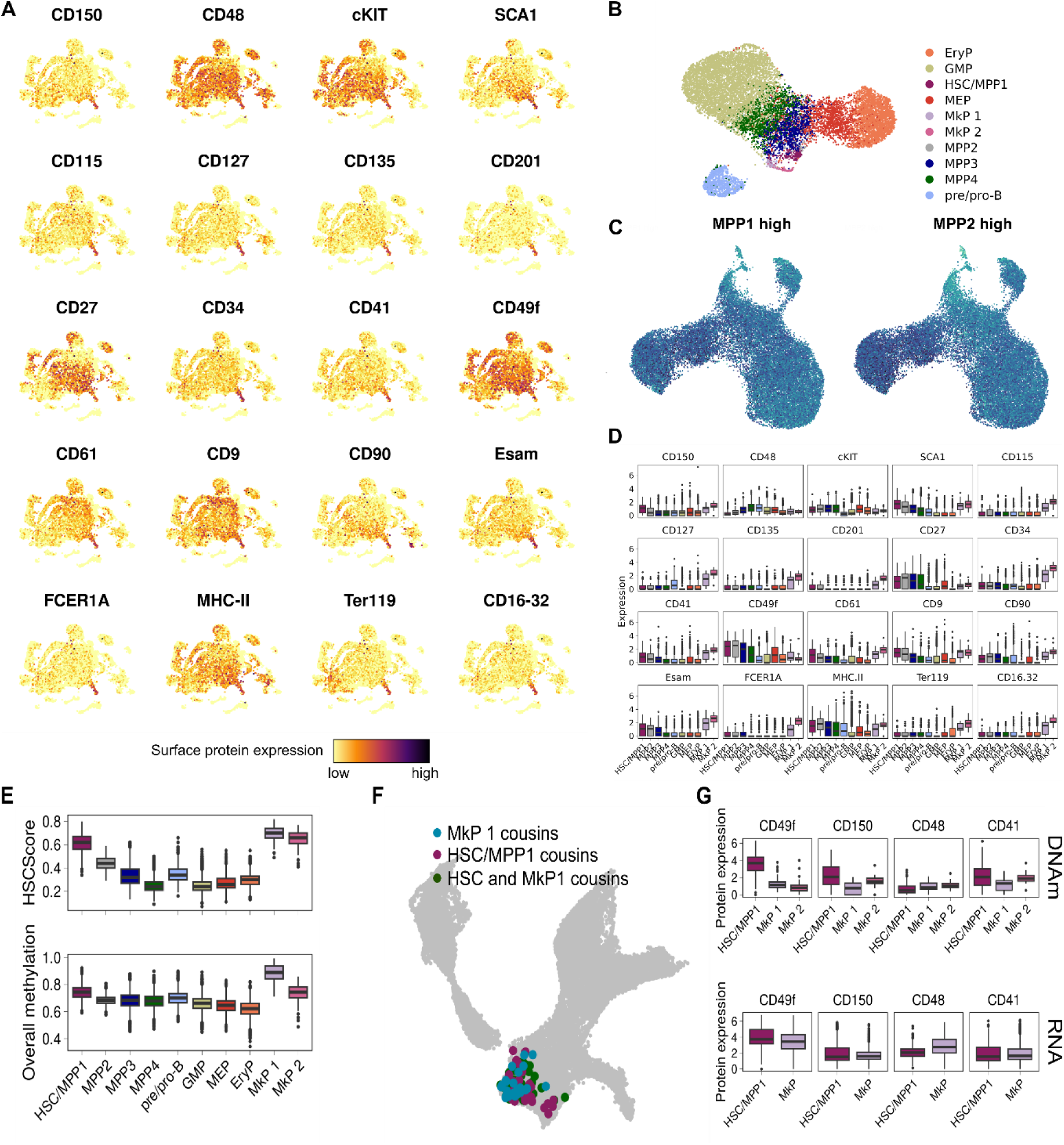
ScTAM-seq delivers a high-resolution map of murine hematopoiesis. **A.** The uMAP from Figure 1C/D is color coded by the expression of 20 surface proteins measured using oligonucleotide-tagged antibodies. **B.** uMAP defined only on the *dynamic* CpGs. The plot shows all 13,885 cells from the main LARRY experiments. Indicated in colors are the cell type labels defined in Figure 2. **C.** Relative methylation state of cells in amplicons specifically methylated in MPP1/MPP2 in bulk data. **D.** Expression of the 20 surface proteins across all the cell types as boxplots. The expression values were normalized using the centered-log-ratio normalization. **E.** HSC score defined on the methylation states of CpGs methylated in HSCs in bulk DNAm data and overall methylation state across all CpGs assayed. **F.** RNA uMAP as in main Figure 1G, highlighting HSC and MkP cells that harbor the same LARRY barcodes observed in the specific DNA methylation clusters HSC and MkP 1. These cells are cousins to the cells classified as HSC/MkP1 in the DNA methylation experiment. **G.** Surface protein expression of the MkP clusters and HSCs in both the DNAm (upper panel) and RNA (lower panel) modality.

**Figure S3:**
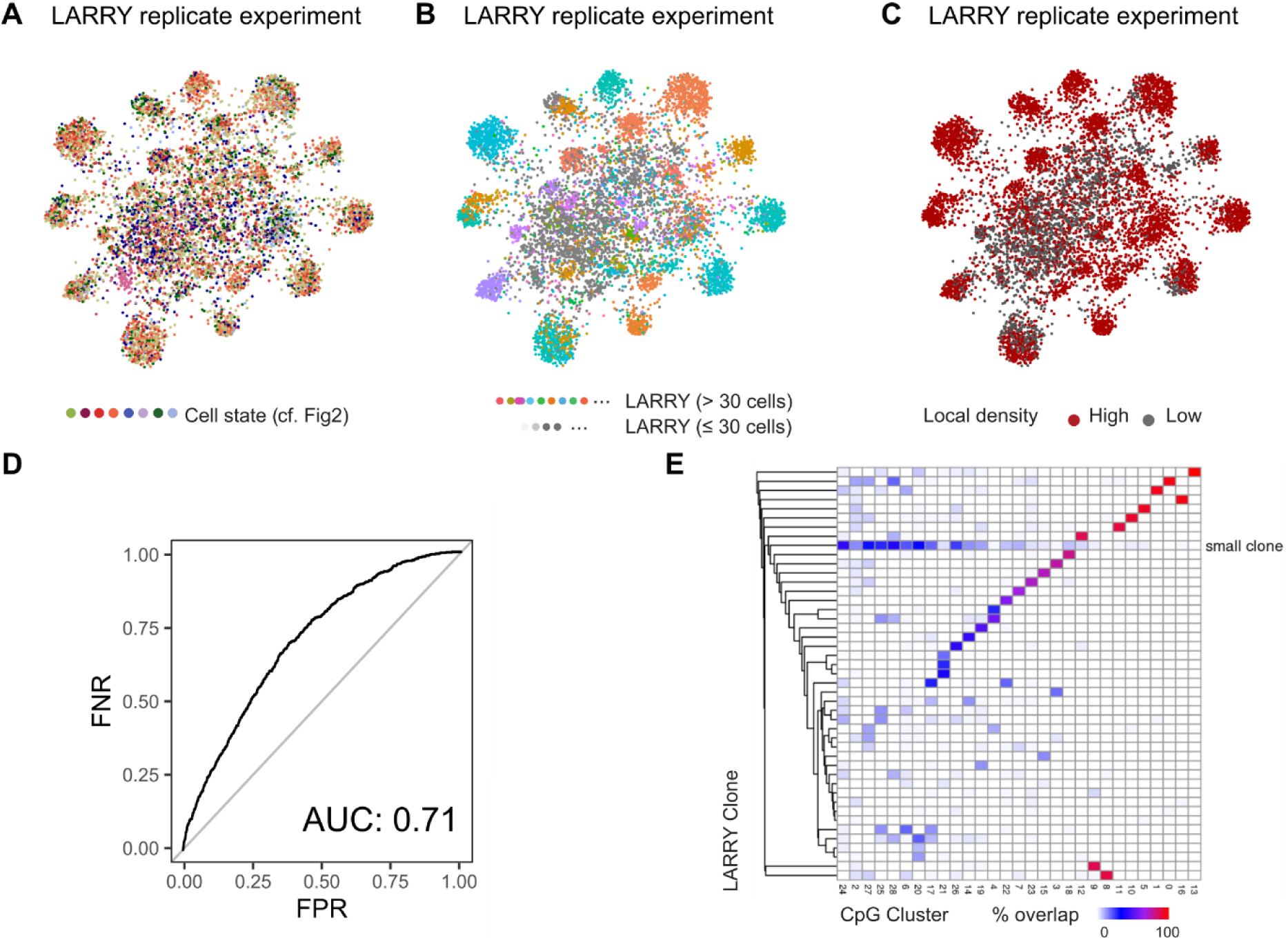
Validation of EPI-clone’s capability on a biological replicate. **A, B, C.** Clonal uMAP as in Figure 3B and C, computed for a biological replicate of the LARRY experiment. Indicated are the cell state (A) and the LARRY barcode (B). C highlights cells that were selected as part of expanded clones, based on local density in PCA space. **D.** Receiver-Operating Characteristics Curve characterizing the performance of the local density criterion in selecting expanded clones for the biological replicate. **E.** Overlap between clones defined using EPI-clone and ground truth labels for the biological replicate.

**Figure S4:**
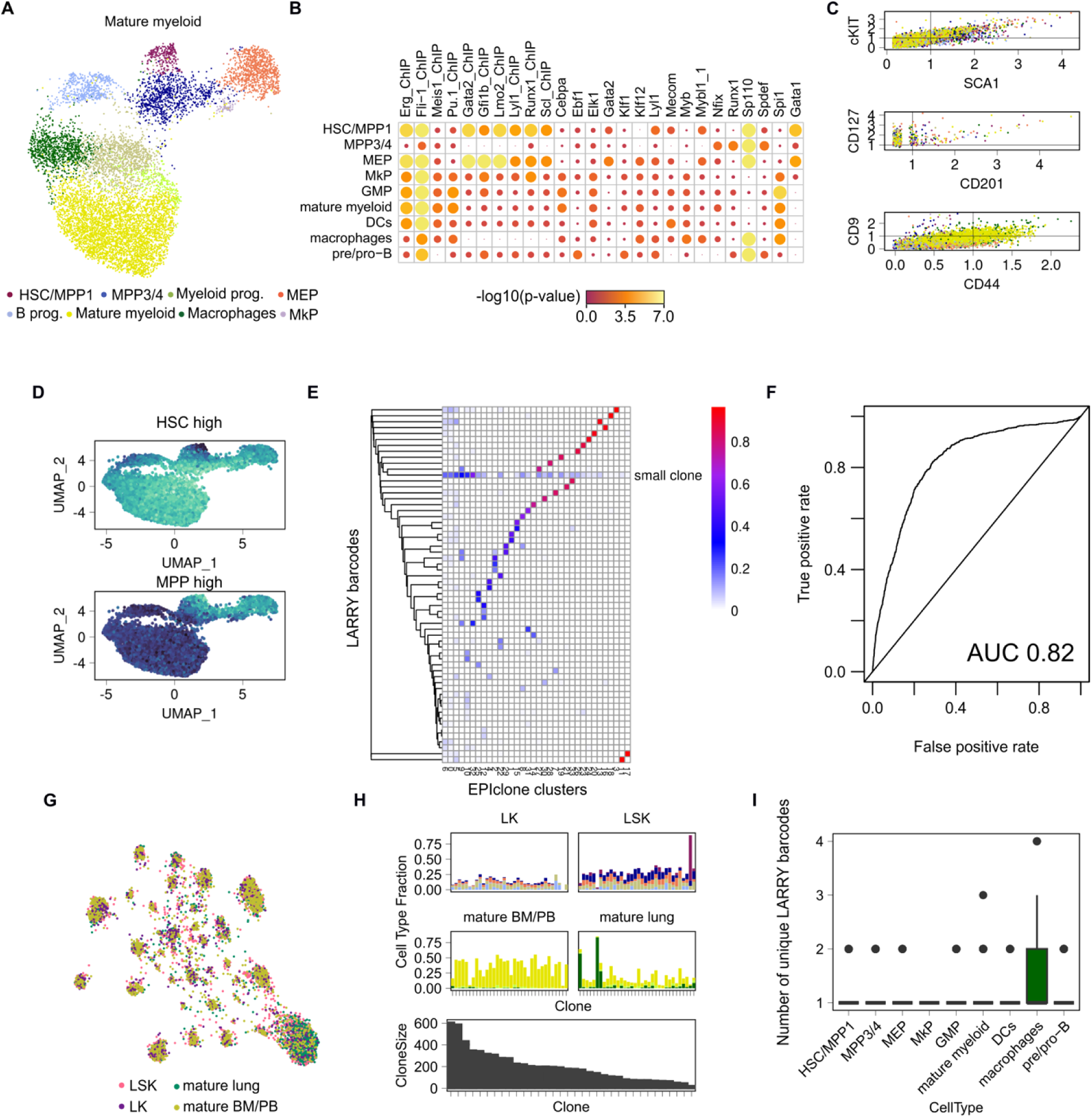
EPI-Clone’s performance in mature myeloid cells. **A.** uMAP based on dynamic CpGs showing the differentiation state of mature myeloid cells and their progenitors. **B.** Enrichment of CpGs specifically unmethylated in a cell-type cluster according to the vicinity to the annotated TFBS. **C.** Expression of surface proteins in the different cell type clusters for stem-cell-specific markers (cKIT, SCA1, CD201) and markers of mature myeloid cells (CD9, CD44). **D**. uMAP as in A, highlighting relative methylation state of cells across all CpGs that are methylated in HSCs or MPP3/4 in bulk data. **E.** Overlap between clones defined using EPI-clone and ground truth clonal labels for the mature myeloid experiment. **F**. Receiver-Operating Characteristics Curve characterizing the performance of the local density criterion in selecting expanded clones for the mature myeloid experiment. **G:** uMAP representation as in Figure 3h visualizing the different cellular compartments including progenitors (LSK, LK) and mature cells from lung and BM/PB. **H.** Cell type distribution and clone sizes in different clones identified by EPI-Clone and stratified by cellular compartment **I.** Number of unique LARRY barcodes per cell type cluster.

**Figure S5:**
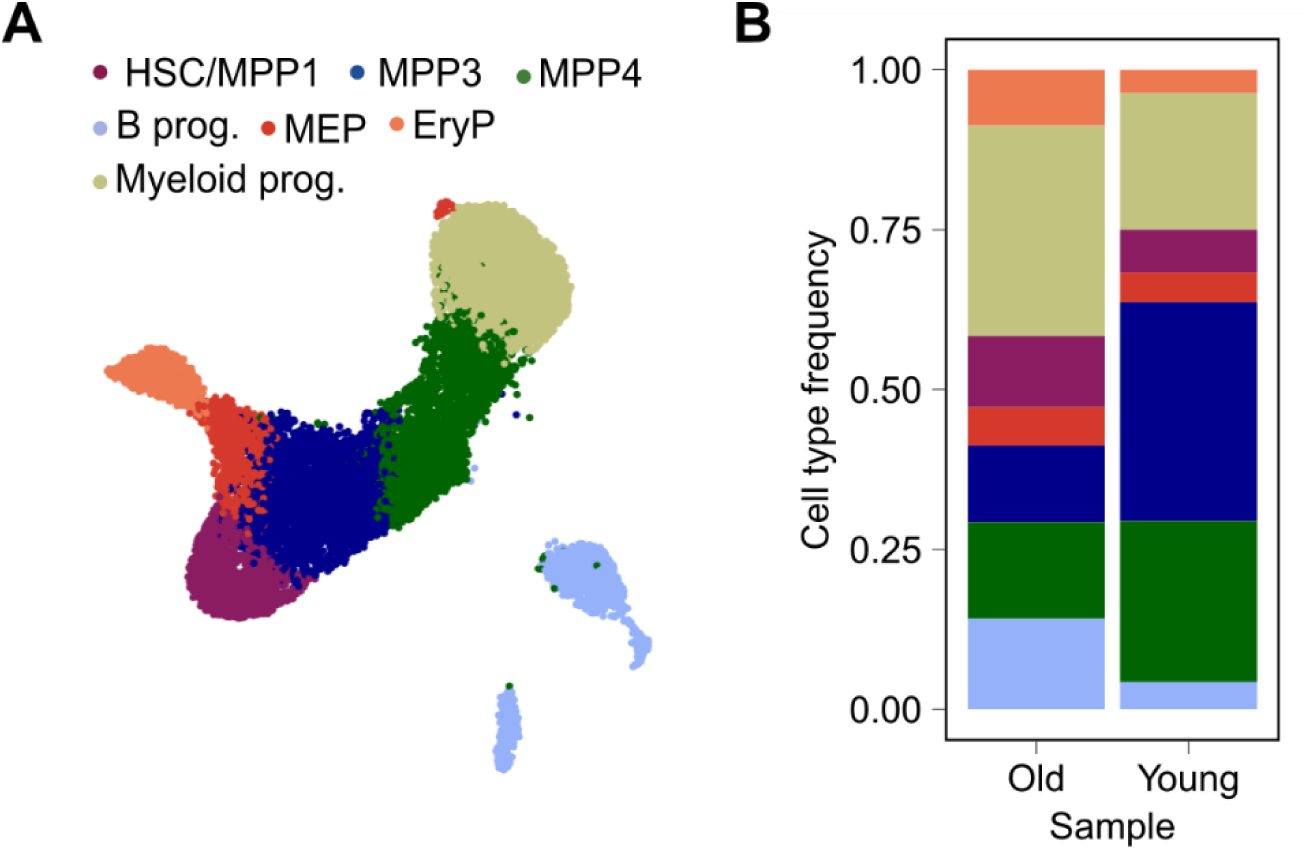
Cell type distributions in the old versus the young mouse. **A.** UMAP based on the dynamic CpGs depicting the cell types identified in the aging experiment (combing young and old mouse). **B.** Comparison of cell type distributions in the old versus the young mouse.

**Figure S6:**
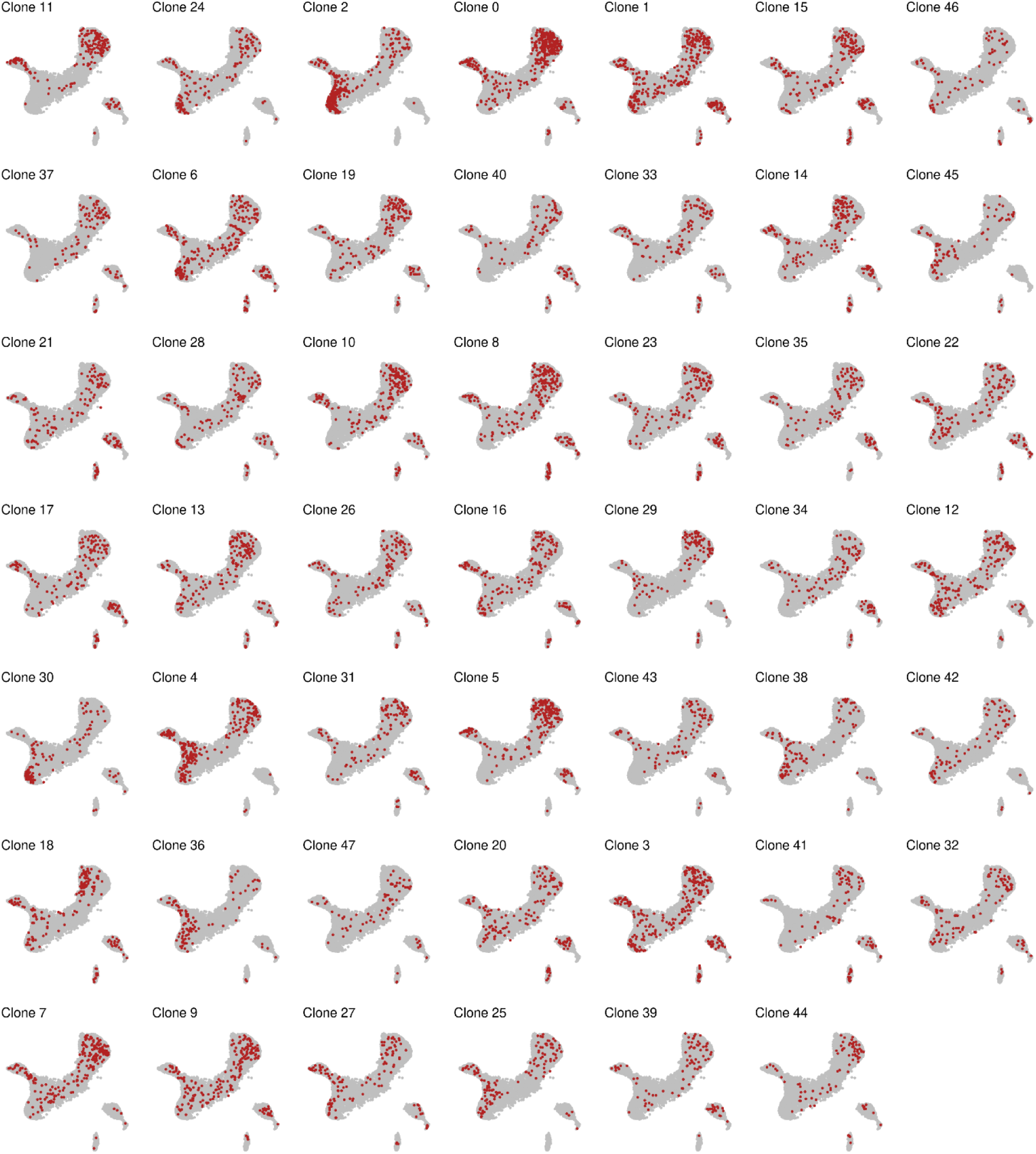
Visualization of clones in the differentiation uMAP for the old mouse.

**Figure S7:**
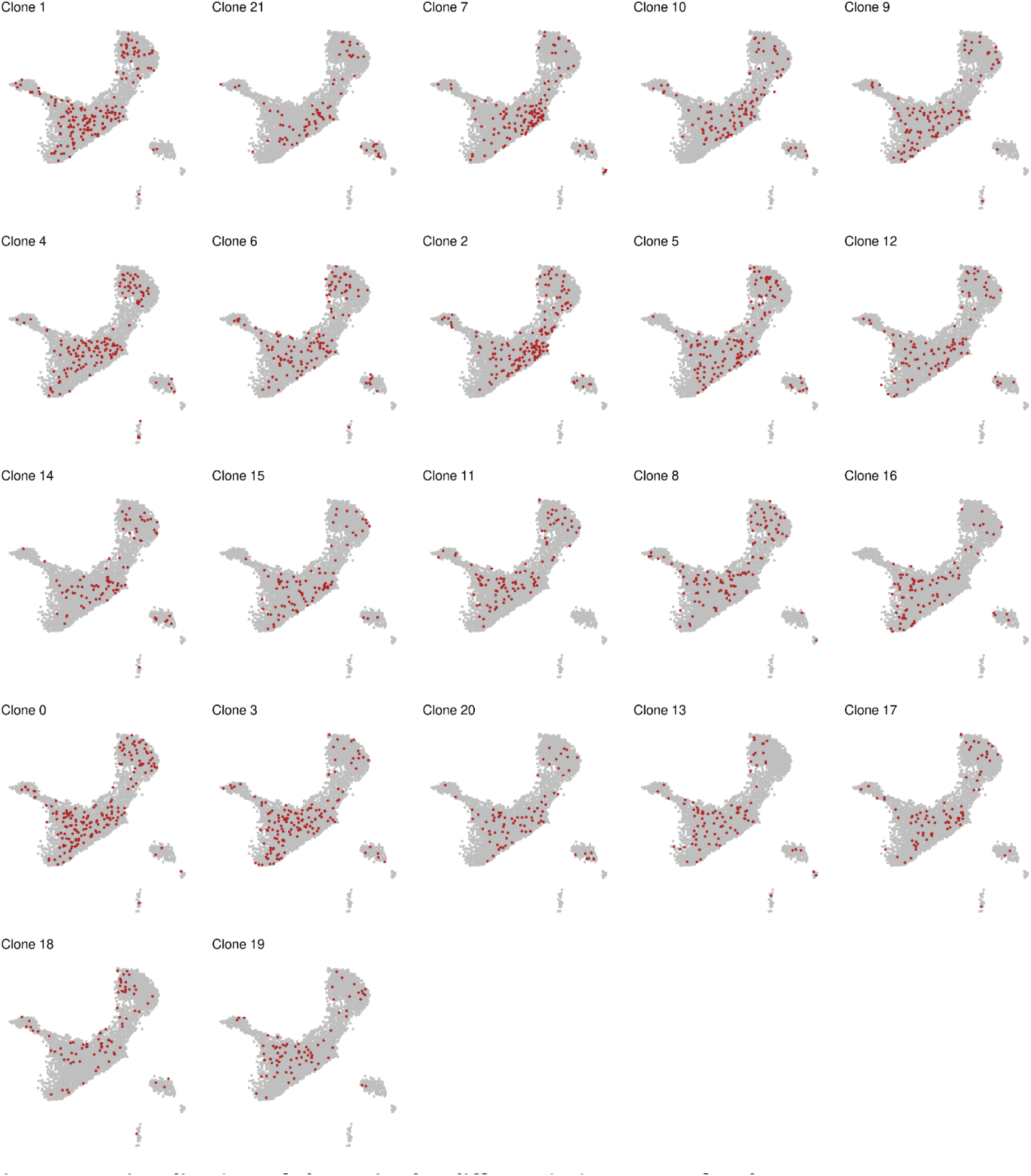
Visualization of clones in the differentiation uMAP for the young mouse.

**Figure S8:**
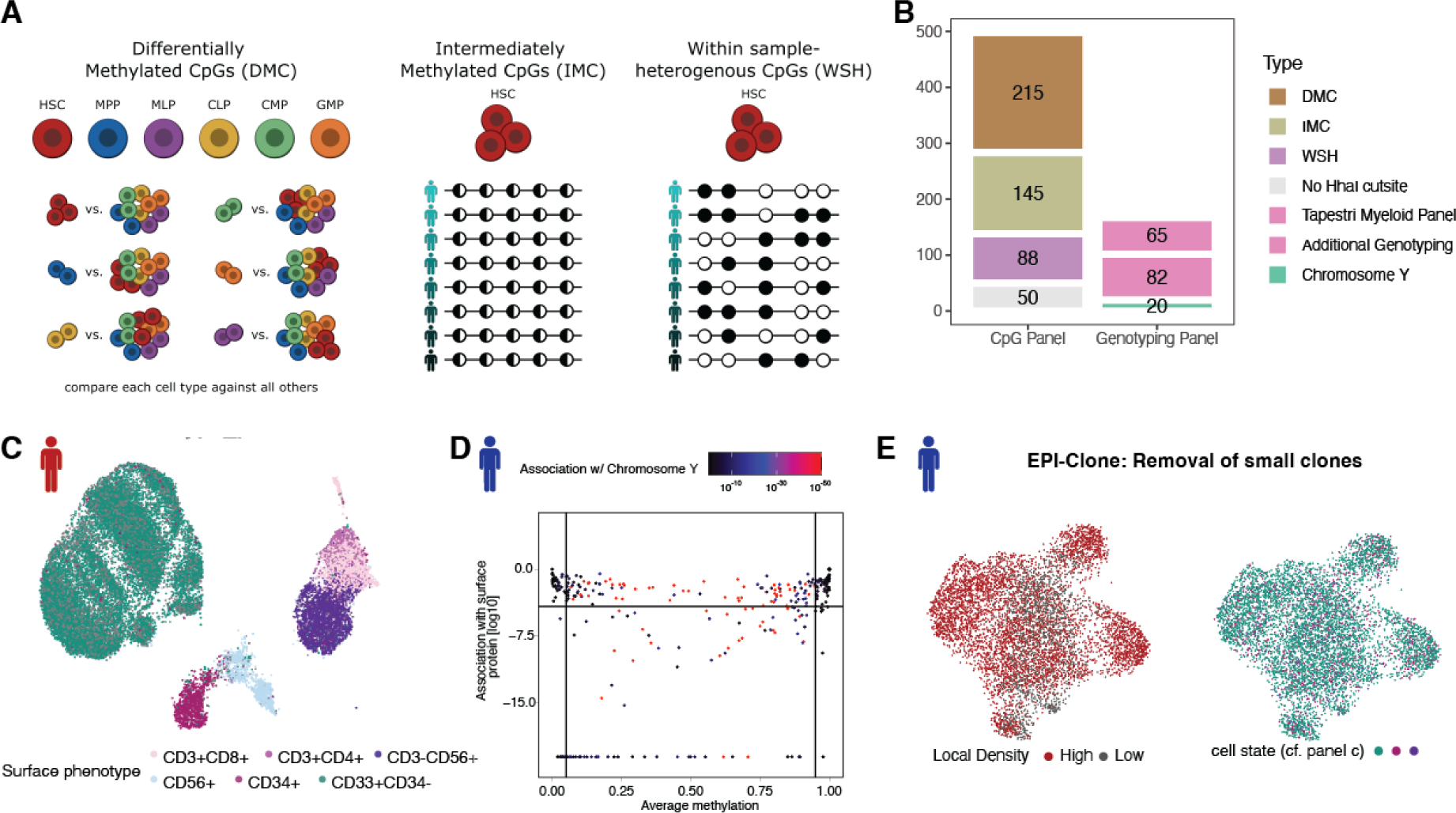
Application of EPI-Clone to human bone marrow samples. **A.** Scheme illustrating selection of target CpGs, see also Methods. **B.** Bar chart illustrating the composition of the panel. **C.** uMAP of all CpGs for donor 2, see also main figure 5b. **D.** Selection of static and dynamic CpGs for donor 1, see also main figure 1e. E. Identification of small clones, see also main figure 3c,d.

**Figure S9:**
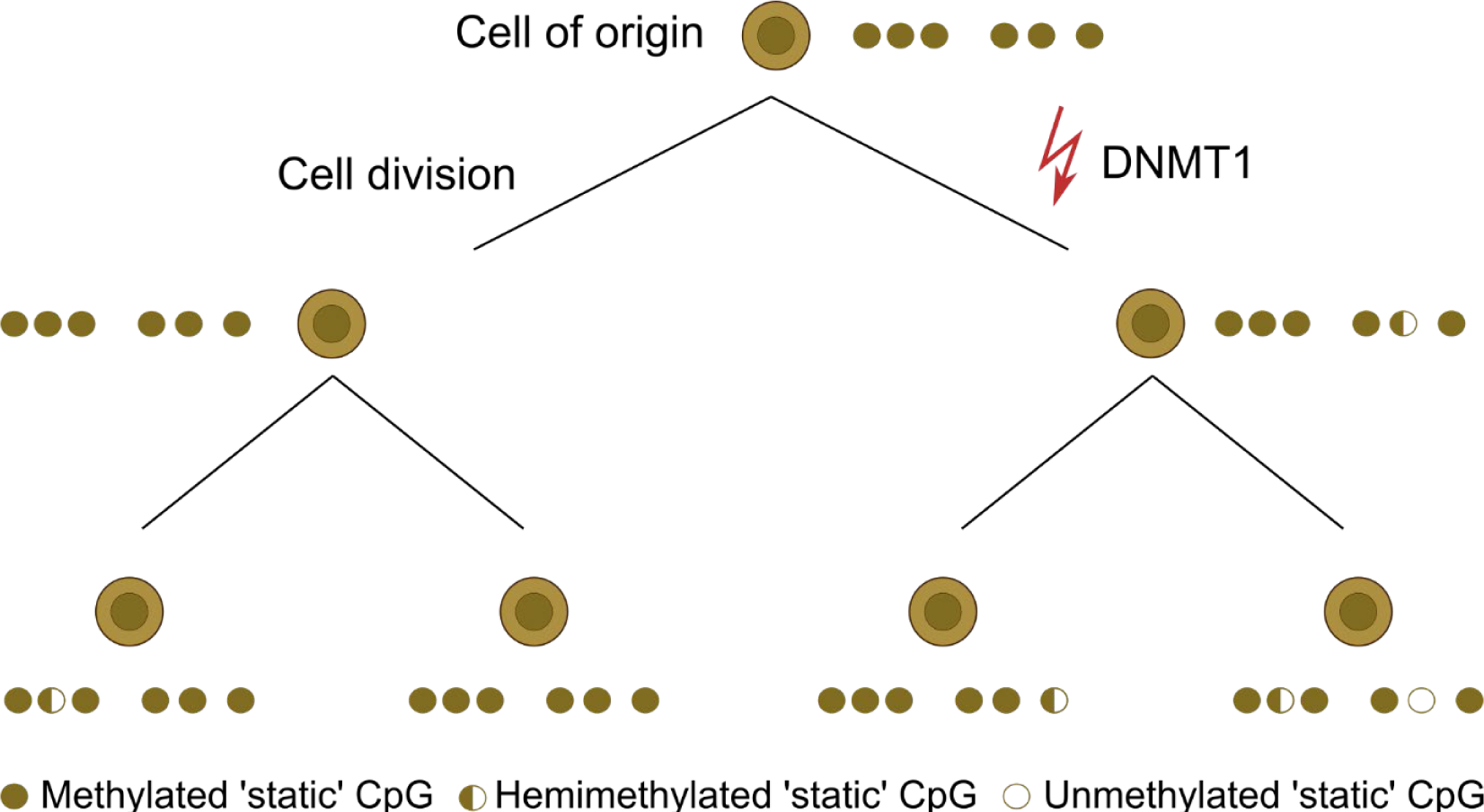
Model for the origin of epimutations. From a fully methylated default state in the cell of origin, epimutations randomly occur as loss of methylation. One potential explanation is the insufficient function of DNMT1 for copying the methylation state to the daughter strand, causing a methylation loss on one of the strands. In a subsequent cellular division, DNA methylation is then lost on both strands.

## Supplementary Tables

Table S1: TotalSeq-B antibodies used for generating surface protein expression libraries.

Table S2: List of primers used for constructing and amplifying LARRY barcodes.

Table S3: List of all amplicons and CpGs covered by the scTAM-seq murine panel with further annotations.

Table S4: Sequencing statistics for scTAM-seq and scRNA-seq libraries.

## Supplementary Note: Characterization of MkP 1 and MkP 2 in single-cell DNA methylation data

Unexpectedly, in DNA methylation data we observed two clearly distinct populations with a surface phenotype of LSK-CD49f-CD150+CD48-CD41+CD61+, consistent with a megakaryocyte progenitor identity (Figure S2D). Interestingly, both these populations resembled HSCs according to bulk methylome profiles (Figure S2E), but differed with regard to their overall methylation level: MkP2 was highly methylated, expressed more CD41, and down-regulated CD49f, relative to HSCs and MkP1 (Figure S2E, S2G). In the scRNA-seq data, no distinct sub-clusters of megakaryocyte precursors were observed. To characterize the gene expression of these putative MkP populations, we made use of the LARRY barcode information that was shared between the RNA-seq and the scTAM-seq experiment.

We observed that MkP1 cells from the methylation experiment had cousins in the scRNAseq experiment that fell into all precursor states, with a particular enrichment of cells near the HSC to megakaryocyte precursor interface (Figure S2F). In particular, these cells were in cycle, lacked expression of myeloid or HSC markers, and instead expressed a large number of genes associated with suppression of HSC proliferation (Apoe, Tgfbr3, Zeb2, Auts2, Ezh2), and also the transcription factors Gata1 and Dach2, as well as the megakaryocyte markers Itga2b at low levels. The close relationship between these cells and the bona fide, Vwf+Itga2b+CD41+CD61+ megakaryocyte progenitors suggests that this population corresponds to a cycling intermediate between HSCs and MkPs. By contrast, MkP2 expressed megakaryocyte markers more highly. Possibly, these cells methylate their DNA extensively to prepare for endomitosis. Notably, we did not observe any overlapping lineage barcodes between MkP2 cells and cell in the scRNA-seq modality.

Together this finding highlights that some cell states may be more visible in the DNAm modality compared to scRNA-seq.

